# Metabolic excretion associated with nutrient-growth dysregulation promotes the rapid evolution of an overt metabolic defect

**DOI:** 10.1101/498543

**Authors:** Robin Green, Sonal, Lin Wang, Samuel F.M. Hart, Wenyun Lu, David Skelding, Justin C. Burton, Hanbing Mi, Aric Capel, Hung Alex Chen, Aaron Lin, Arvind R. Subramaniam, Joshua D. Rabinowitz, Wenying Shou

## Abstract

In eukaryotes, conserved mechanisms ensure that cell growth is coordinated with nutrient availability. Overactive growth during nutrient limitation (“nutrient-growth dysregulation”) can lead to rapid cell death. Here, we demonstrate that cells can adapt to nutrient-growth dysregulation by evolving major metabolic defects. Specifically, when yeast lysine auxotrophic mutant *lys*^*-*^ encountered lysine limitation, an evolutionarily novel stress, cells suffered nutrient-growth dysregulation. A sub-population repeatedly evolved to lose the ability to synthesize organosulfurs (*lys*^*-*^*orgS*^*-*^). Organosulfurs, mainly glutathione and glutathione conjugates, were released by *lys*^*-*^ cells during lysine limitation when growth was dysregulated, but not during glucose limitation when growth was regulated. Limiting organosulfurs conferred a frequency-dependent fitness advantage to *lys*^*-*^*orgS*^*-*^ by eliciting a proper slow growth program including autophagy. Thus, nutrient-growth dysregulation is associated with rapid organosulfur release, which enables the selection of organosulfur auxotrophy to better tune cell growth to the metabolic environment. We speculate that evolutionarily novel stresses can trigger atypical release of certain metabolites, setting the stage for the evolution of new ecological interactions.

## Introduction

All organisms must coordinate growth with the availability of nutrients. In eukaryotes, when nutrients are abundant, cells express growth-promoting genes and grow. When nutrients are limited, cells halt growth and instead launch stress-response programs to survive. Mechanisms that ensure this coordination between nutrient availability and growth are conserved across eukaryotes (Jewell and Guan, 2013).

The budding yeast *S. cerevisiae* senses and responds to the availability of natural nutrients, nutrients that must be supplied from the environment (Zaman et al., 2008). Examples of natural nutrients include carbon, nitrogen, phosphorus, and sulfur. When natural nutrients are abundant, the TORC1 (target of rapamycin complex 1) pathway is activated. If the carbon source happens to be glucose, the Ras/protein kinase A (PKA) pathway is additionally activated (Zaman et al., 2009, 2008). Activated TORC1 and PKA pathways promote growth-related processes, including ribosome synthesis, biomass accumulation, and cell division (Fig 1A green box). Simultaneously, TORC1 and PKA inhibit stress-responsive processes (Fig 1A red box). Thus, abundant natural nutrients set cell state to the growth mode. In contrast, when one of the essential nutrients is missing, cell state is switched to the stress-response mode (Fig 1A&B red box): Cells upregulate stress-responsive genes, and acquire enhanced resistance to heat and to high osmolarity. Cell division is arrested in an unbudded state; oxidative metabolism is elevated, wherein cells consume more oxygen, and do not ferment glucose into ethanol (Brauer et al., 2008; Klosinska et al., 2011; Petti et al., 2011; Saldanha et al., 2004). Additionally, cells engage in autophagy, a stress survival process involving degradation and recycling of cytosol and organelles (Huang and Klionsky, 2007; Klosinska et al., 2011; Slavov and Botstein, 2011). Thus, proper nutrient-growth regulation allows cells to grow when natural nutrients are abundant, and to maintain high viability when natural nutrients are scarce or lacking.

**Fig 1.**
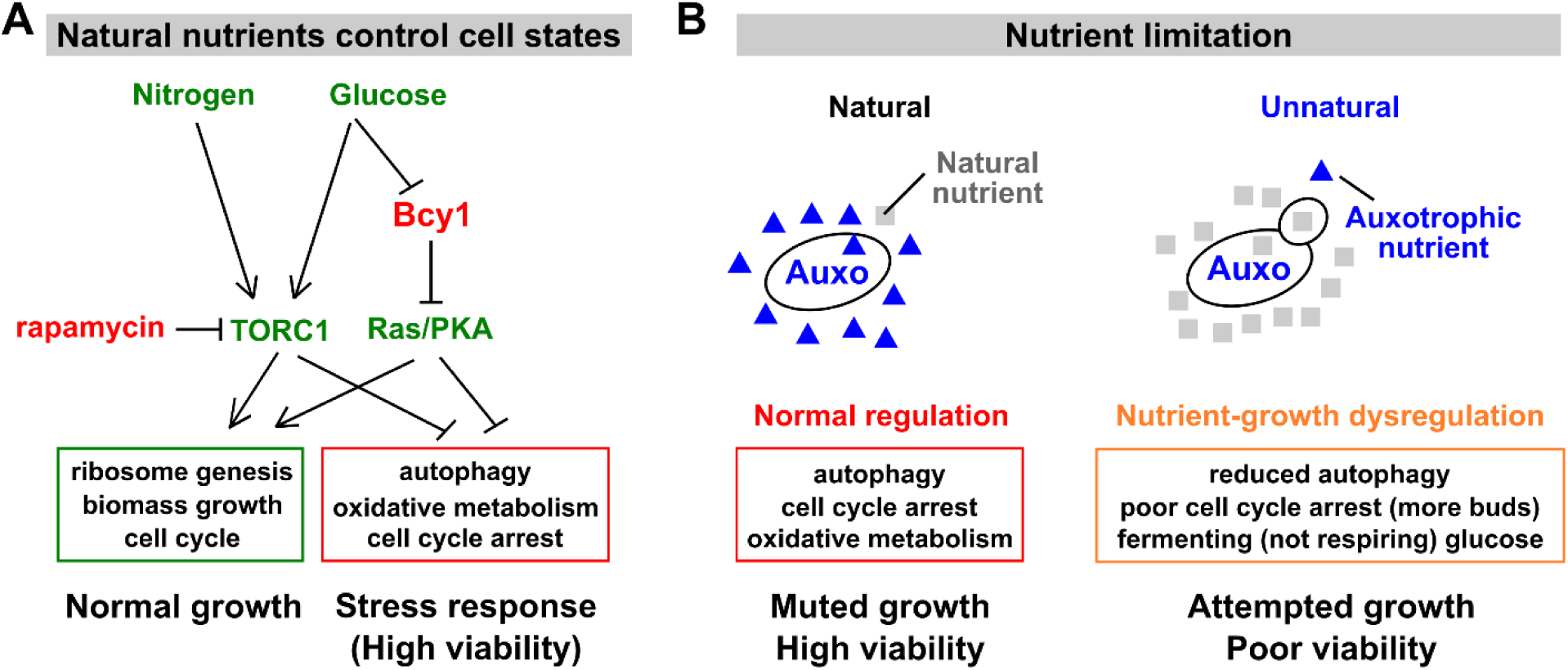
Nutrient-growth regulation and dysregulation. (**A**) Natural nutrients control the growth or stress state of a cell (reviewed in (Zaman et al., 2008)). Growth stimulatory molecules are colored green, and growth inhibitory molecules are colored red. Broadly speaking, the presence of natural essential nutrients (e.g. nitrogen, glucose, sulfur, phosphorus) activate the TORC1 pathway. Glucose additionally activates the Ras/PKA pathway, although this activation is transient if essential natural nutrients are incomplete (Thevelein et al., 2000). For simplicity, we have omitted other input pathways (e.g. Conrad et al., 2014). TORC1 and Ras/PKA pathways activate cell growth (green box) and inhibit stress response (red box). Conversely, the shortage of a natural nutrient or inhibiting TORC1 via rapamycin triggers stress responses including cell cycle arrest, autophagy, and oxidative metabolism. Note that the mRNA levels of greater than a quarter of yeast genes are linearly correlated with growth rate, independent of the nature of the nutrient (Brauer et al., 2008; Regenberg et al., 2006). During slow growth, repressed genes include those involved in ribosome synthesis, translation initiation, and protein and RNA metabolism, while induced genes are involved in autophagy, lipid metabolism, and oxidative metabolism (including those annotated to peroxisomes and the peroxisomal matrix) (Brauer et al., 2008; Castrillo et al., 2007; Gasch et al., 2000; Slavov and Botstein, 2011). (**B**) Left: When limited for natural nutrient, an auxotroph responds properly (red box) and survives with high viability. Right: When limited for the auxotrophic nutrient, an auxotroph suffers nutrient-growth dysregulation (orange box): Despite nutrient limitation, cells experience poor cell cycle arrest and reduced autophagy, and metabolize glucose via fermentation instead of respiration. Consequently, these cells suffer low viability. Supplementary Text 1 offers additional discussions.

In contrast, “unnatural limitation” occurs when, for example, a yeast auxotrophic mutant is limited for the metabolite it can no longer make (e.g. a *leu-* mutant limited for leucine). Auxotrophic limitation is unusual in nature: Because wild yeast strains are diploid, they remain prototrophic (capable of synthesizing a metabolite) even if the functionality of one of the two gene copies is impaired. However, auxotrophic limitation might mimic natural limitation in some cases. For example, intracellular glutamine level may reflect nitrogen availability (Boer et al., 2010), while intracellular methionine level may reflect sulfur or general amino acid availability (Laxman et al., 2014; Sutter et al., 2013). Leucine, whose biosynthesis occurs in both cytoplasm and mitochondria, may monitor the tricarboxylic acid (TCA) cycle and the ADP:ATP ratio (Kingsbury et al., 2015). Thus, some of the mechanisms for sensing and responding to these amino acids (González and Hall, 2017; Zhang et al., 2018) might actually function to sense natural nutrients.

Nutrient-growth regulation can become dysfunctional during unnatural auxotrophic limitation. During unnatural limitation, cells fail to arrest cell cycle, metabolize glucose through fermentation instead of respiration, and suppress the transcription of stress-response genes, vacuolar and autophagy genes, as well as respiration-related genes involved in mitochondria and the TCA cycle (Brauer et al., 2008; Petti et al., 2011; Saldanha et al., 2004; Slavov and Botstein, 2013) (Fig 1B, orange box). Cells engage in significantly less autophagic activity under unnatural auxotrophic starvation compared to natural nitrogen starvation (Table II in (Takeshige et al., 1992)). Unnaturally starved cells suffer poor viability which can be rescued by inhibiting growth (Boer et al., 2008; Gresham et al., 2011; Petti et al., 2011), a phenotype we define as “nutrient-growth dysregulation”.

The model of nutrient-growth regulation and dysregulation during nutrient scarcity (Fig 1B, left and right panels, respectively) is supported by multiple experiments. Inactivating the TORC1 pathway, either by using its inhibitor rapamycin or by deleting *TOR1* or its downstream effector *SCH9*, allowed auxotrophic cells to better survive unnatural limitations (Boer et al., 2008; Gresham et al., 2011). Deleting *PPM1*, a gene that promotes growth and inhibits autophagy, aided *leu*^*-*^ cells to survive unnatural leucine starvation, but not natural phosphate starvation (Boer et al., 2008; Gresham et al., 2011). Deletion of *BCY1*, which resulted in constitutively active PKA, caused failed cell cycle arrest and poor viability during nitrogen starvation (Matsumoto et al., 1983; Toda et al., 1987). Finally, auxotrophic cells under unnatural limitation died much faster when the carbon source was glucose (which additionally activates PKA) compared to when the carbon source was poor such as ethanol or glycerol (Boer et al., 2008).

Since nutrient-growth regulation is conserved across eukaryotes (Jewell and Guan, 2013), here we investigate how cells suffering nutrient-growth dysregulation might evolve, taking advantage of the *lys2Δ* strain (“*lys*^*-*^”) of *S. cerevisiae*.

## Results

### Nutrient-growth dysregulation in *lys*^*-*^ cells under lysine limitation

Previous work has established that *lys*^*-*^ cells limited for lysine displayed hallmarks of nutrient-growth dysregulation (Slavov and Botstein, 2013). Unlike prototrophic cells limited for a natural nutrient, lysine-limited *lys*^*-*^ cells suffered decoupling between the cell division cycle and biomass production, fermented glucose despite abundant oxygen, and suppressed the transcription of autophagy genes and mitochondrial and respiration genes (Slavov and Botstein, 2013). Note that oxidative metabolism during nutrient limitation is important: Deleting genes involved in mitochondrion organization and respiration rendered cells sensitive to glucose starvation (Klosinska et al., 2011).

Consistent with nutrient-growth dysregulation, lysine-limited *lys*^*-*^ cells suffer low viability that can be rescued by simultaneously inhibiting cell growth. We used a lysine auxotrophic *S. cerevisiae* strain *lys*^*-*^, engineered to harbor a *lys2* deletion mutation and also expressing the fluorescent protein mCherry (WY1335/WY2490, Table S1). When starved for lysine, *lys*^*-*^ cells rapidly lost fluorescence (Fig 2A, blue circles), indicative of rapid cell death (Fig S1). Rapid death of *lys-* cells during lysine starvation was prevented by rapamycin – an inhibitor of the growth activator TORC1 (Fig 2A, green crosses), or by removing the growth activator glucose (Fig 2A, green squares). The viability of lysine-starved *lys-* cells was further reduced by deleting *BCY1*, an inhibitor of the Ras/PKA growth pathway (Fig 2A, red diamonds; Fig S2; Fig 1A).

**Fig 2.**
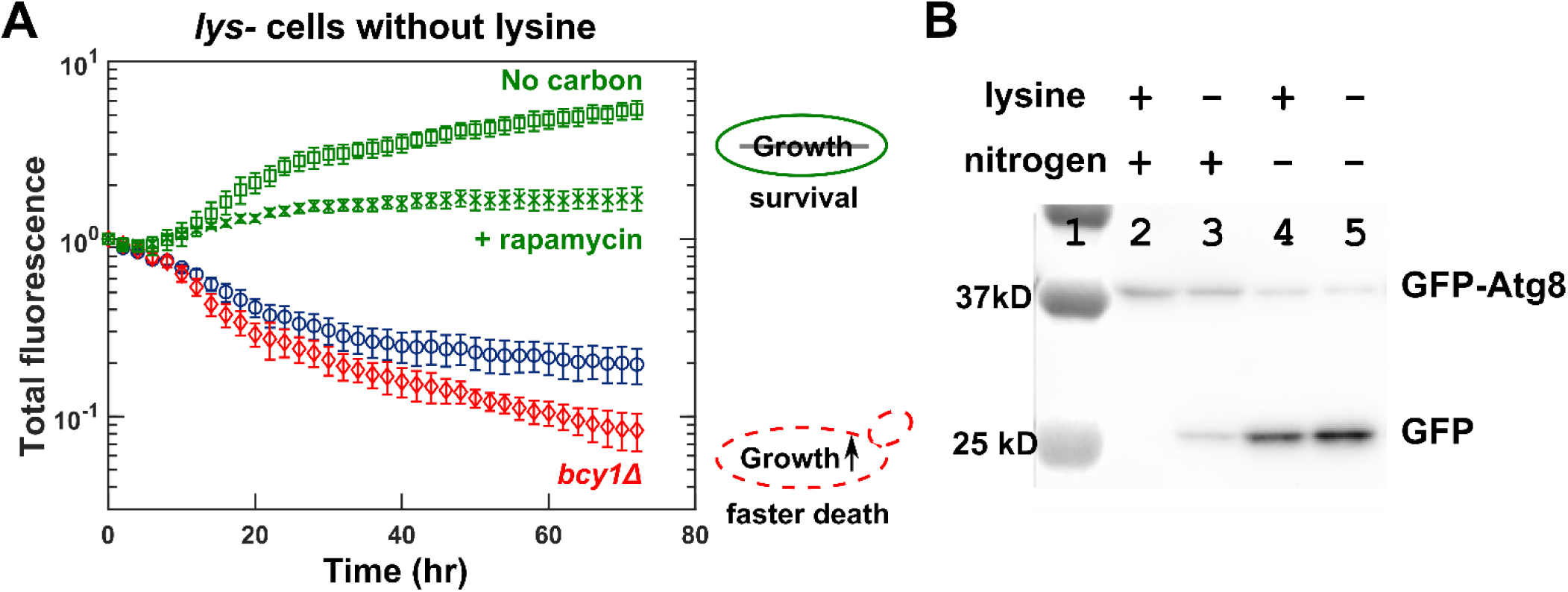
*lys*^*-*^ cells suffer nutrient-growth dysregulation when limited for lysine. **(A) The poor viability of *lys*^-^cells during lysine starvation is rescued by inhibiting growth and is exacerbated by activating growth.** Exponentially growing mCherry-expressing *lys*^-^(WY2490) cells were washed and starved for either only lysine or both lysine and glucose for 3 hours to deplete cellular storage, and cultured and imaged in indicated environments (“Fluorescence microscopy”). Total fluorescence (normalized to the initial value) approximates total biomass (Hart et al., 2019b). In the absence of lysine, *lys*^-^cells died rapidly when glucose was abundant (2%, blue circles). This rapid death could be rescued if we inhibited growth by adding the TORC1 inhibitor rapamycin (1 µM, green crosses) or by simultaneously starving for glucose (“No carbon”, green squares). Rapid death was exacerbated if we activated growth by deleting the PKA inhibitor *BCY1* (red diamonds). Error bars correspond to two standard deviations for 6 replicate wells. (B) ***lys*^-^cells engage in less autophagy during lysine starvation compared to during nitrogen starvation**. Lane 1 is the protein ladder. *lys*^-^(WY2521) cells were grown to the exponential phase (Lane 2) and washed free of nutrients. Cells were starved for only lysine (Lane 3), only nitrogen (Lane 4), or both lysine and nitrogen (Lane 5) for 8 hrs. Cell extracts were subjected to Western blotting using anti-GFP antibodies (Methods, “Autophagy assay”). High GFP:GFP-Atg8 ratio indicates high autophagy activity.

Consistent with nutrient-growth dysregulation, autophagy is reduced in *lys*^*-*^ cells during lysine limitation compared to natural limitation. We used a GFP-Atg8 cleavage assay (Xie et al., 2008) to monitor autophagy. Specifically, when cells undergo autophagy, GFP-Atg8 is delivered to the vacuole and cleaved, and the free GFP has high resistance to vacuolar degradation. Thus, the ratio of GFP to GFP-Atg8 is a metric for autophagy activity (Torggler et al., 2017). *lys*^*-*^ cells starved for lysine showed significantly less autophagy activity compared to *lys-* cells deprived of the natural nutrient nitrogen or simultaneously starved for nitrogen and lysine (Fig 2B).

In summary, *lys*^*-*^ cells limited for lysine suffer nutrient-growth dysregulation manifested as reduced viability and autophagy compared to during natural limitation.

### Repeated evolution of organosulfur auxotrophy

To examine how cells might cope with nutrient-growth dysregulation, we evolved *lys*^*-*^ cells in lysine limitation for tens of generations, either as monocultures in lysine-limited chemostats or in cocultures with a lysine-releasing strain (Methods, “Evolution”; “Chemostats and turbidostats”). We then randomly chose a total of ∼70 evolved *lys*^*-*^ clones for whole-genome sequencing. Similar to our previous findings (Hart et al., 2019a; Waite and Shou, 2012), each clone carried at least one mutation that increased the cell’s affinity for lysine. These mutations included duplication of Chromosome 14 which harbors the high-affinity lysine permease gene *LYP1* (Hart & Pineda et al., 2019), or losing or reducing activities of genes involved in degrading Lyp1 (e.g. *ECM21, RSP5*, and *DOA4*) (Lin et al., 2008; Waite and Shou, 2012) (Table S2).

Surprisingly, even though the input medium contained no organosulfurs (sulfur-containing organic molecules; blue in Fig 3B), ∼10% of the ∼70 sequenced clones harbored mutations in the biosynthetic pathway that converts externally supplied sulfate to essential organosulfurs. These mutations included *met10, met14*, and *met17*, and arose from independent chemostat and coculture lines (Fig 3A box). Indeed, these mutant clones required an external source of organosulfur such as methionine to grow (Fig 3A box) (Masselot and Robichon-Szulmajster, 1975). We designate these mutants as *lys*^*-*^*orgS*^*-*^. *lys*^*-*^*orgS*^*-*^ arose after the initial evolutionary adaptation to lysine limitation, since *lys*^*-*^ and *lys*^*-*^*orgS*^*-*^ clones from the same culture harbored an identical *ecm21* or *rsp5* mutation (Table S2, matched color shading). We also found a glutamine auxotroph *gln1* (Fig 3A, see Discussions). Since we observed *gln1* only once, we focused on the evolution of *orgS*^*-*^.

**Fig 3.**
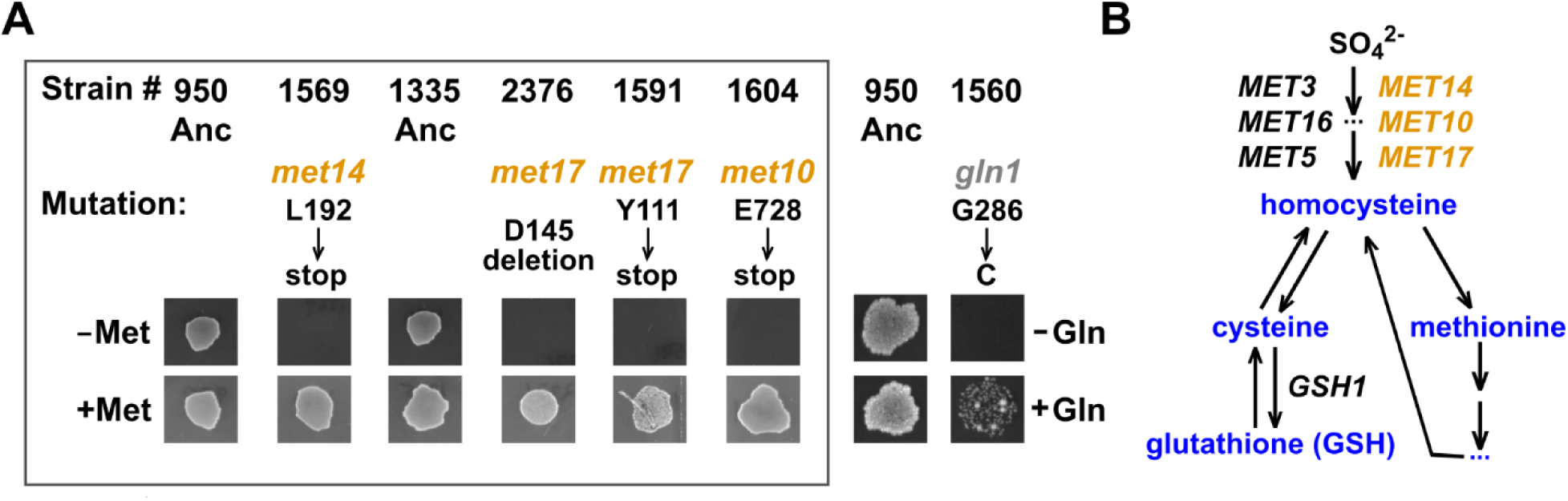
The evolution of auxotrophy. (**A**) **The evolution of auxotrophs**. Ancestral *lys-* cells (WY950; WY1335) were grown for tens of generations in minimal medium, either in lysine-limited chemostats (Novick and Szilard, 1951) or via co-culturing with a lysine releaser in a cross-feeding yeast community (Hart et al., 2019a; Shou et al., 2007). Out of 20 independent lines, we randomly isolated ∼70 clones for whole-genome sequencing. Chemostat evolution and coculture evolution both yielded *met-* mutants: 1 (WY1604) out of 9 clones in chemostat evolution; 3 (WY1569, WY2376, WY1591) out of ∼60 clones in coculture evolution). These mutants, all isolated from independent lines, required an externally supplied organosulfur such as methionine to grow. A glutamine auxotrophic *gln1* clone was also identified. In the experiment shown here, clones were grown to exponential phase in SD supplemented with amino acids, washed with SD, starved for 3 hours to deplete cellular storage, and spotted on indicated agar plates at 30°C. (**B**) **The organosulfur synthesis pathway in *S. cerevisiae***. *S. cerevisiae* utilizes sulfate supplied in the medium to synthesize the organosulfur homocysteine, which is then used to make a variety of other organosulfurs including methionine, cysteine, and glutathione (Thomas and Surdin-Kerjan, 1997). All *lys*^-^*orgS*^-^mutants we have identified (orange) fail to synthesize homocysteine, and thus can be supported by any organosulfurs depicted here (blue) which are inter-convertible.

When we tested six independently-evolved cultures (three chemostat monocultures and three cross-feeding cocultures) at 30∼80 generations (using the growth assay in Fig 3A), *lys*^*-*^*orgS*^*-*^ mutants could be detected in all cases (Fig S3; Methods “Quantifying auxotroph frequency”), suggesting repeated evolution of organosulfur auxotrophy. Below, we focus on *lys*^*-*^ monocultures since in these cultures, evolved *lys*^*-*^*orgS*^*-*^ cells must have received organosulfurs from *lys*^*-*^ cells.

### The organosulfur niche mainly consists of glutathione S-conjugates and glutathione

We initially tested whether chemostat supernatant contained methionine or cysteine using gas chromatography. Both compounds were undetectable. We then resorted to liquid chromatography-mass spectrometry (Methods, “LC-MS”). We identified glutathione (GSH) as a released organosulfur (Fig 4A, S4 and S5). GSH, a tri-peptide comprising glutamate, cysteine, and glycine, is a major cellular redox buffer, and can form glutathione-S-conjugates (GSX) with itself (GS-SG) and with other compounds via disulfide bonds (e.g. GS-S-CoA) (Ishikawa, 1992). Indeed, *gsh1*^*-*^ cells, whose growth can be supported by externally supplied GSH and GS-SG but not methionine (Fig S5B) (Grant et al., 1996), grew in chemostat supernatants (Fig S5E). GSH release rate was comparable between ancestral and evolved *lys-* cells in lysine-limited chemostats (Fig S7D) and thus, organosulfur supply was uninterrupted during evolution.

**Fig 4.**
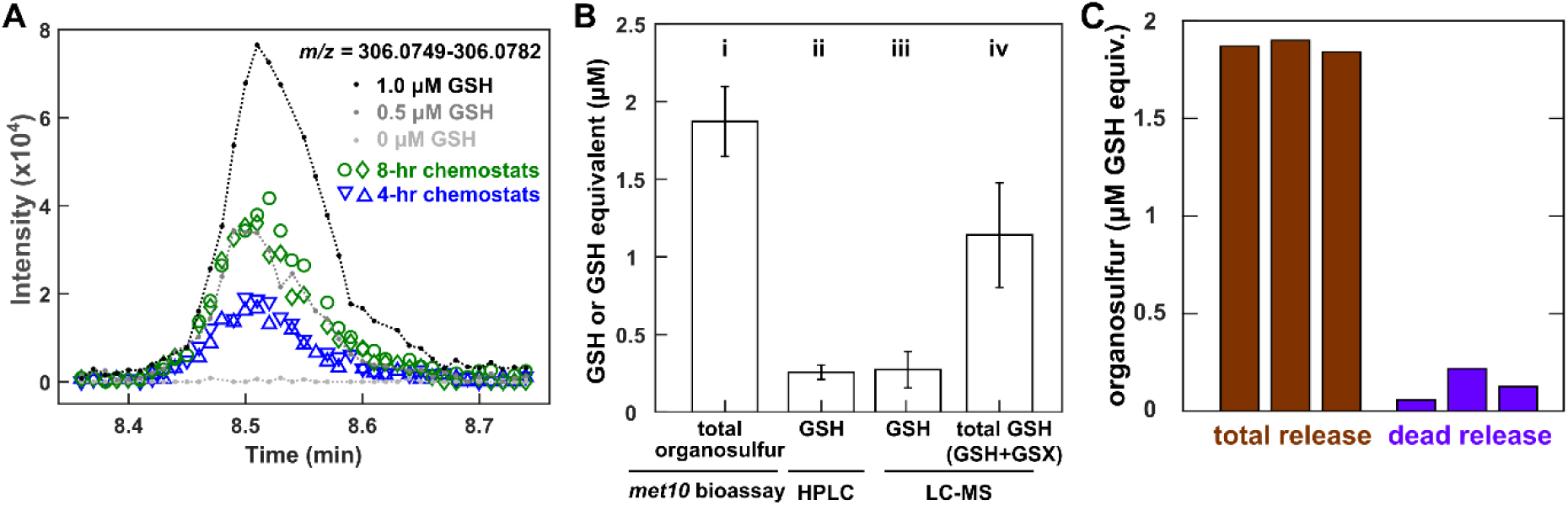
Lysine-limited *lys-* cells mainly release glutathione and glutathione S-conjugates. Ancestral *lys-* cells (WY1335) were cultured in lysine-limited chemostats at 8-hr doubling time unless otherwise indicated. (**A**) **Mass spectra traces of reduced glutathione (GSH) ion**. Gray and black lines correspond to known quantities of GSH in growth media. Blue and green correspond to filtered supernatants harvested at ∼48 hrs. (**B**) **GSH and GSH S-conjugates (GSX) constitute the majority of the organosulfur niche**. Supernatants were harvested at steady state (∼26 hr) and filtered. (**i**) Bioassay quantification of total organosulfur was performed by comparing the final turbidity of *met10*- grown in supernatants versus in various known concentrations of GSH. (**ii, iii**) GSH in supernatant was quantified by HPLC and LC-MS. (**iv**) GSH+GSX in supernatants were quantified by reducing GSX to GSH with TCEP and measuring total GSH via LC-MS. Error bars mark two standard deviations of samples from three independent chemostats. (**C**) **Organosulfurs are likely released by live cells**. Total organosulfur in chemostat supernatant (brown) far exceeded that expected from release by dead cells (purple). Organosulfurs were quantified using the *met10* bioassay. To quantify dead cell release, we measured dead cell density using flow cytometry (Methods, “Flow cytometry”), and multiplied it with the average amount of organosulfur per cell (Methods, “Metabolite extraction”). Three independent experiments are plotted.

To estimate the total organosulfur niche in chemostat supernatant, we developed a yield-based bioassay (Methods, “Bioassays”). We used *met10*^*-*^, a mutant isolated in our evolution experiment which can use a variety of organosulfurs including GSH, GS-SG, and methionine (Fig S5C, D). By comparing the final turbidity yield of *met10*^*-*^ in a chemostat supernatant versus in minimal medium supplemented with various known amounts of GSH, we can estimate supernatant organosulfurs in terms of GSH equivalents (Fig 4B, i). GSH quantified via HPLC and LC-MS (Methods) was much lower than the total organosulfur niche (Fig 4B, compare ii and iii with i). We then chemically reduced chemostat supernatants to convert GSX to GSH, and measured total GSH via LC-MS (Methods). We found that GSH and GSX together dominated the organosulfur niche (Fig 4B, compare i and iv), with GSX being the major component (Fig 4B, compare iii with iv).

The organosulfur niche was mainly created by live cell release rather than by dead cell lysis. Specifically, we compared the organosulfur concentration quantified in chemostat supernatant (Fig 4C, brown) with that expected from dead cell release (calculated by multiplying dead cell density with average intracellular organosulfur content; Methods “Metabolite extraction”) (Fig 4C, purple). Our measurements suggest that organosulfurs were mainly released by live cells (Fig 4C; Fig S6), consistent with the observation that GSH and GSX are exported from the cell in an ATP-dependent fashion (Ishikawa, 1992; Rebbeor et al., 1998).

### Organosulfur release is associated with nutrient-growth dysregulation, not slow growth

To test whether organosulfur release is associated with nutrient-growth dysregulation, we grew *lys*^*-*^ cells in turbidostats (Gresham and Dunham, 2014) (Methods, “Chemostats and turbidostats”) where excess lysine supported fast growth and thus nutrient-growth regulation should be normal. *lys*^*-*^ cells in excess lysine released GSH at a significantly slower rate than in lysine-limited chemostats (Fig S7, A-C).

To further test whether organosulfur release is associated with nutrient-growth dysregulation or with slow growth, we compared *lys*^*-*^ cells in glucose-limited chemostats versus lysine-limited chemostats at the same doubling time. Nutrient-growth regulation should be normal during glucose limitation but not during lysine limitation (Fig 1B). Indeed, the percent dead cells was much lower during glucose limitation than during lysine limitation (Fig 5A, note the logarithm scale). Organosulfurs accumulated to a steady state in lysine-limited chemostats, but were undetectable in glucose-limited chemostats, even though live population density was higher in glucose-limited chemostats (Fig 5A-B).

**Fig 5.**
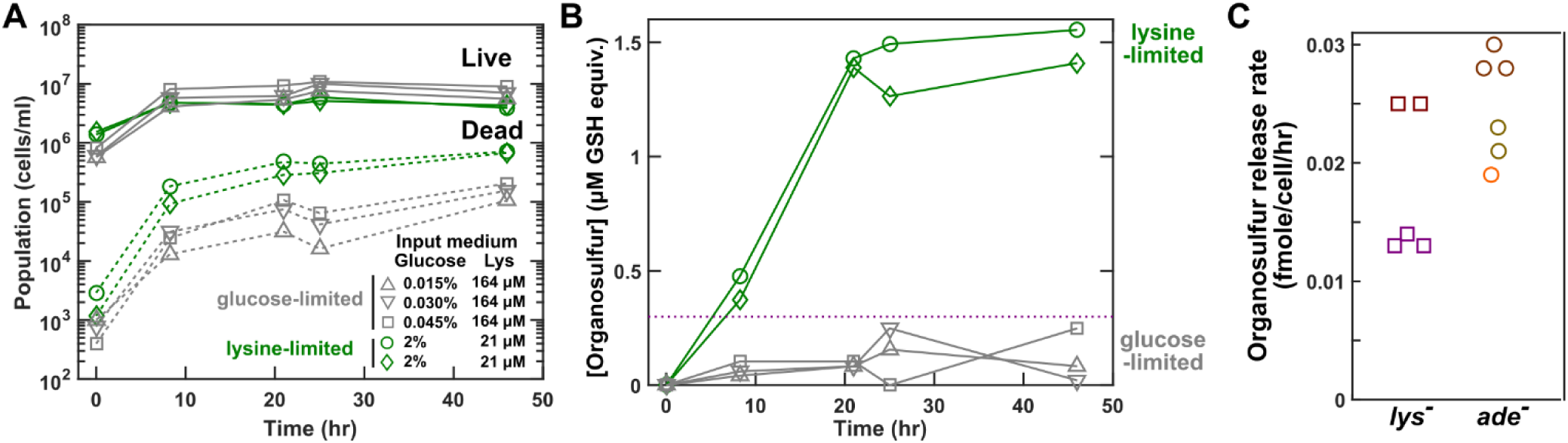
Organosulfur release is associated with nutrient-growth dysregulation. *lys-* cells (WY1335) were cultured in lysine-limited (green) or glucose-limited (grey) chemostats at 8-hr doubling time. The concentrations of glucose and lysine in the input fresh minimal medium are marked. (**A**) **Higher percent dead cells in lysine-limited chemostats than in glucose-limited chemostats**. Live and dead cell densities were measured via flow cytometry (Methods, “Flow cytometry”). (**B**) **Higher organosulfur concentrations in lysine-limited chemostats than in glucose-limited chemostats**. Organosulfur concentration was measured in terms of GSH equivalents using the *met10* bioassay (Fig S5), with brown dotted line marking the lower limit of the linear detection range. (**C**) **Adenine-limited *ade***^***-***^ **cells release organosulfurs at comparable rates as lysine-limited *lys***^***-***^ **cells**. Cells were cultured in chemostats limited for the respective auxotrophic metabolite at 8-hr doubling time until a steady state has been reached. Release rate was quantified using *r* = *dil* * [orgS]_*ss*_/[Live]_*ss*_ (Eq. 14 in (Hart et al., 2019a)) where *dil* is the dilution rate (*ln*2/8 /hr), [orgS]_*ss*_ is the steady state organosulfur concentration (measured in terms of GSH equivalents using the *met10* bioassay; Fig S5), and [Live]_*ss*_ is the steady state live cell density. Different colors correspond to experiments done on different days, and each symbol represents an independent chemostat.

Organosulfurs are also released by nutrient-growth dysregulated *ade*^*-*^ cells in adenine-limited chemostats. Adenine-limited *ade*^*-*^ cells released organosulfurs at a similar rate as lysine-limited *lys*^*-*^ cells (Fig 5C), and this release was mediated by live cells (Fig S8). Although differing from lysine-limited *lys*^*-*^ cells in several aspects (Fig S9), adenine-limited *ade*^*-*^ cells displayed nutrient-growth dysregulation: *ade*^*-*^ cells lost viability during adenine starvation, and this low viability was rescued by removing glucose (Fig S9).

Taken together, these results suggest that organosulfur release is associated with nutrient-growth dysregulation, not with slow growth.

### Organosulfur limitation confers a frequency-dependent fitness advantage to *orgS*^*-*^

*lys*^*-*^*orgS*^*-*^ repeatedly rose from one mutant cell to a detectable frequency in independent cultures. This suggests that the *orgS*^*-*^ mutation may confer a fitness benefit to *lys-* cells. To test this, we randomly chose an evolved *lys*^*-*^*orgS*^*-*^ clone (WY1604), restored its *orgS*^*-*^ (*met10*) mutation to wild type, and compared isogenic *lys*^*-*^*orgS*^*-*^ and *lys*^*-*^ clones in a variety of nutrient environments.

According to the prevalent “energy saving” hypothesis (D’Souza et al., 2014; Zamenhof and Eichhorn, 1967), excess organosulfurs should help *lys*^*-*^*orgS*^*-*^ by sparing its cost of *de novo* synthesis of organosulfurs. However, we observed the opposite: when both GSH and lysine were in excess, *lys*^*-*^ *orgS*^*-*^ grew significantly slower than *lys*^*-*^ (Fig 6A). In excess GSH but limiting lysine, *lys*^*-*^*orgS*^*-*^ behaved similarly to *lys*^*-*^, surviving poorly but regaining viability when growth was inhibited by rapamycin (Fig 6B Right, compare orange with blue; Fig S10B).

**Fig 6.**
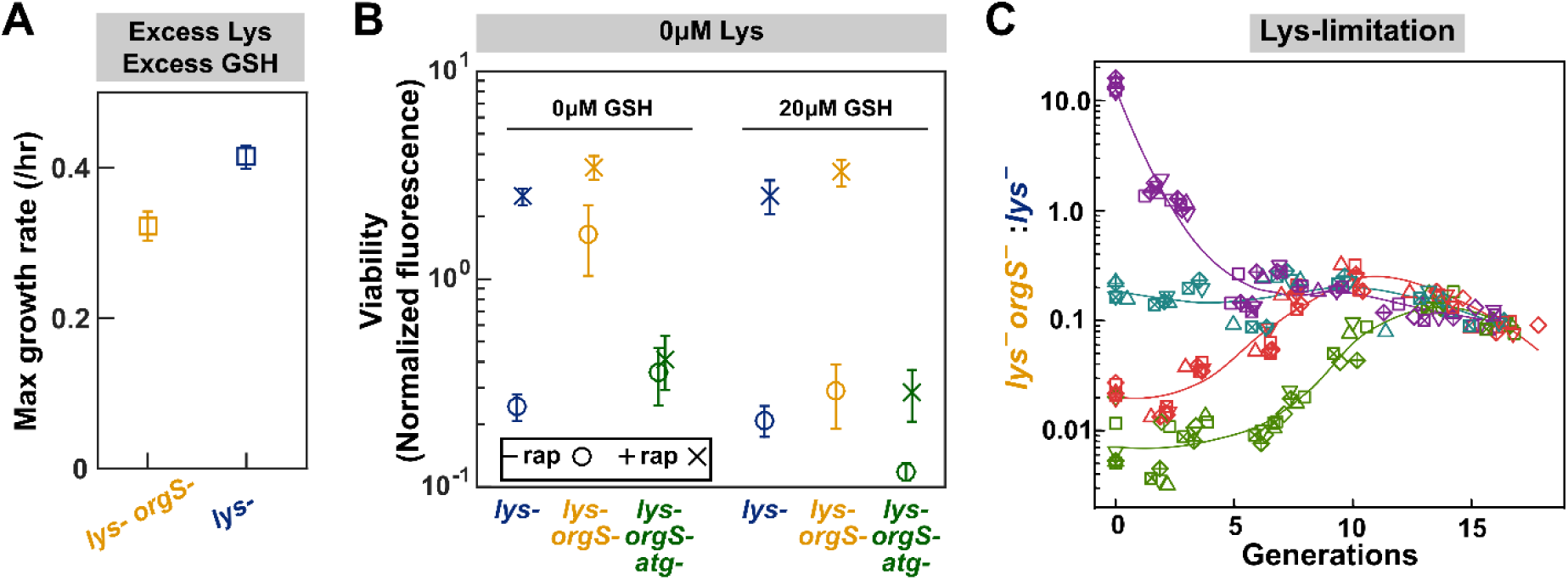
*lys*^-^*orgS*^-^displays a frequency-dependent fitness advantage over *lys*^-^during lysine limitation. (**A, B**) **Fitness advantage of *lys*^-^*orgS*^-^over *lys*^-^requires low organosulfur and autophagy**. Cell growth was imaged via fluorescence microscopy. (**A**) In excess lysine and GSH, *lys*^-^*orgS*^-^(WY1604, orange) grew slower than isogenic *lys*^-^(WY2429, blue). (**B**) Isogenic *lys*^-^(WY2429, blue), *lys*^-^*orgS*^-^(WY1604, orange), and *lys*^-^*orgS*^-^*atg5*^-^ (WY 2370, green) were cultured in the absence of lysine without (circles) or with (crosses) 1 µM rapamycin and without or with 20 µM GSH. Total fluorescence of the field of view at the endpoint (110 hrs) was normalized against that of time zero. *lys*^-^*orgS*^-^(yellow circle, 2^nd^ column) survived lysine starvation better than *lys*^-^(blue circle, 1^st^ column) when GSH was low. When GSH was abundant, *lys*^-^*orgS*^-^behaved similarly to *lys*^-^, surviving lysine starvation poorly (yellow circle, 5^th^ column) and rescued by the growth inhibitor rapamycin (yellow cross, 5^th^ column). The high viability of organosulfur-limited *lys*^-^*orgS*^-^and of rapamycin-treated cells were abolished when autophagy was prevented (WY2370, green circles and crosses). Full data are plotted in Fig S10. Error bars represent two SEM (standard error of mean) from six wells. (**C**) **Negative frequency-dependent fitness advantage of *lys*^-^*orgS*^-^over *lys*^-^**. Isogenic BFP-tagged *lys*^-^*orgS*^-^(WY2072 or WY2073) and mCherry-tagged *lys*^-^(WY2045) were competed in a lysine-limited environment by coculturing with a lysine-releasing strain (WY1340) (Methods). The ratio of *lys*^-^*orgS*^-^to *lys*^-^over time was measured by flow cytometry (Methods, “Competition”). Smooth lines serve as visual guide. Different colors represent different starting ratios, while different symbols represent independent experiments.

We then tested an alternative hypothesis: Unable to process inorganic sulfur in the medium but able to utilize organosulfurs released by *lys*^*-*^, *lys*^*-*^*orgS*^*-*^ cells may respond to this organosulfur limitation similarly as to a natural limitation, mounting a stress response including autophagy. This might in turn confer *lys*^*-*^*orgS*^*-*^ a fitness advantage over nutrient-growth dysregulated *lys*^*-*^ cells. Indeed, when organosulfur was limiting, *lys*^*-*^*orgS*^*-*^ survived lysine limitation better than *lys*^*-*^ (Fig 6B Left, orange circle higher than blue circle; Fig S10A). A similar trend was observed at intermediate levels of lysine and GSH (Fig S11). We would ideally like to compare autophagy activities between *lys*^*-*^*orgS*^*-*^ and *lys*^*-*^ cells, but there are multiple technical challenges (see Fig S12 legend for an explanation). In a less ideal comparison, *lys*^*-*^*orgS*^*-*^ showed lower autophagy activity during lysine starvation than during either organosulfur starvation or organosulfur/lysine double starvation (Fig S12). Importantly, deletion of *ATG5*, a gene essential for autophagy, diminished the advantage of *lys*^*-*^*orgS*^*-*^ over *lys*^*-*^ during lysine limitation (Fig 6B, orange symbols higher than green symbols; Fig S10).

To directly test any fitness difference between isogenic *lys*^*-*^ and *lys*^*-*^*orgS*^*-*^ clones, we marked them with different fluorescent proteins, and competed them in lysine limitation (Methods. “Competition”). When *lys*^*-*^*orgS*^*-*^ was initially abundant (Fig 6C purple), its frequency declined as expected due to competition for the limited organosulfurs released from rare *lys*^*-*^. When *lys*^*-*^*orgS*^*-*^ was rare, its frequency initially increased (Fig 6C red and green), demonstrating a fitness advantage over *lys*^*-*^. Regardless of the starting point, the ratio of *lys*^*-*^*orgS*^*-*^ to *lys*^*-*^ converged to a steady state value (Fig 6C). The observed steady state ratio is consistent with organosulfur release and consumption measurements: At steady state species ratio, organosulfur releasers and consumers must grow at an identical rate, and organosulfur concentration is at a steady state. Assume that all organosulfur compounds are consumable by *lys*^*-*^*orgS*^*-*^ as GSH equivalents. The organosulfur concentration *O* at steady state can be described as 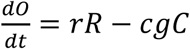, where *R* and *C* are the concentrations of releaser and consumer cells, respectively, *r* represents the release rate of organosulfurs per releaser cell, *g* represents growth rate (= dilution rate in chemostats), and *c* represents the amount of organosulfur consumed per consumer birth. Organosulfur release rate by *lys*^*-*^ in 8-hr chemostats (*g*=ln2/8/hr) was *r*∼0.02 fmole GSH equivalent/cell/hr (Fig 5C). From the slope of the organosulfur-OD standard curve (e.g. Fig S5 blue curves; 1 OD∼ 3×10^7^ cells/ml in our setup), organosulfur consumption per birth *c* ∼2 fmole GSH/cell. Setting the above equation to zero, we obtain 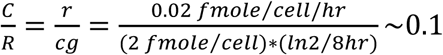, consistent with the observed ratio of ∼0.1 (Fig 6C).

In summary, by helping to at least partially restore nutrient-growth regulation, organosulfur limitation confers rare *lys*^*-*^*orgS*^*-*^ a frequency-dependent fitness advantage over *lys*^*-*^ during lysine limitation (Fig S13).

## Discussions

### Nutrient-growth dysregulation and the evolution of metabolic dependence

Our work demonstrates that within tens of generations, lysine-limited *lys*^*-*^ cells often evolved into two subpopulations *lys*^*-*^ and *lys*^*-*^*orgS*^*-*^ (Fig 3 and Fig S3). *lys*^*-*^ cells initially adapted to lysine limitation via mutations such as *ecm21, rsp5*, and *DISOMY14*. Increased lysine permease Lyp1 on the cell membrane enabled these mutants to outcompete ancestral cells in lysine-limited environments (Hart & Pineda et al., 2019; Hart et al., 2019a; Waite and Shou, 2012).

After this initial adaptation to lysine limitation, two aspects of nutrient-growth dysregulation encouraged the evolution of *orgS*^*-*^ (Fig 7). First, lysine-limited *lys*^*-*^ cells released organosulfurs consisting primarily of GSX and, to a lesser extent, GSH (Figs 4, S4, S6, and S7). This created an organosulfur niche for the survival of *lys*^*-*^*orgS*^*-*^ (Fig S5). Second, when *lys*^*-*^*orgS*^*-*^ initially arose, the mutant enjoyed a fitness advantage over *lys*^*-*^ (Fig 6C and Fig S13).

**Fig 7.**
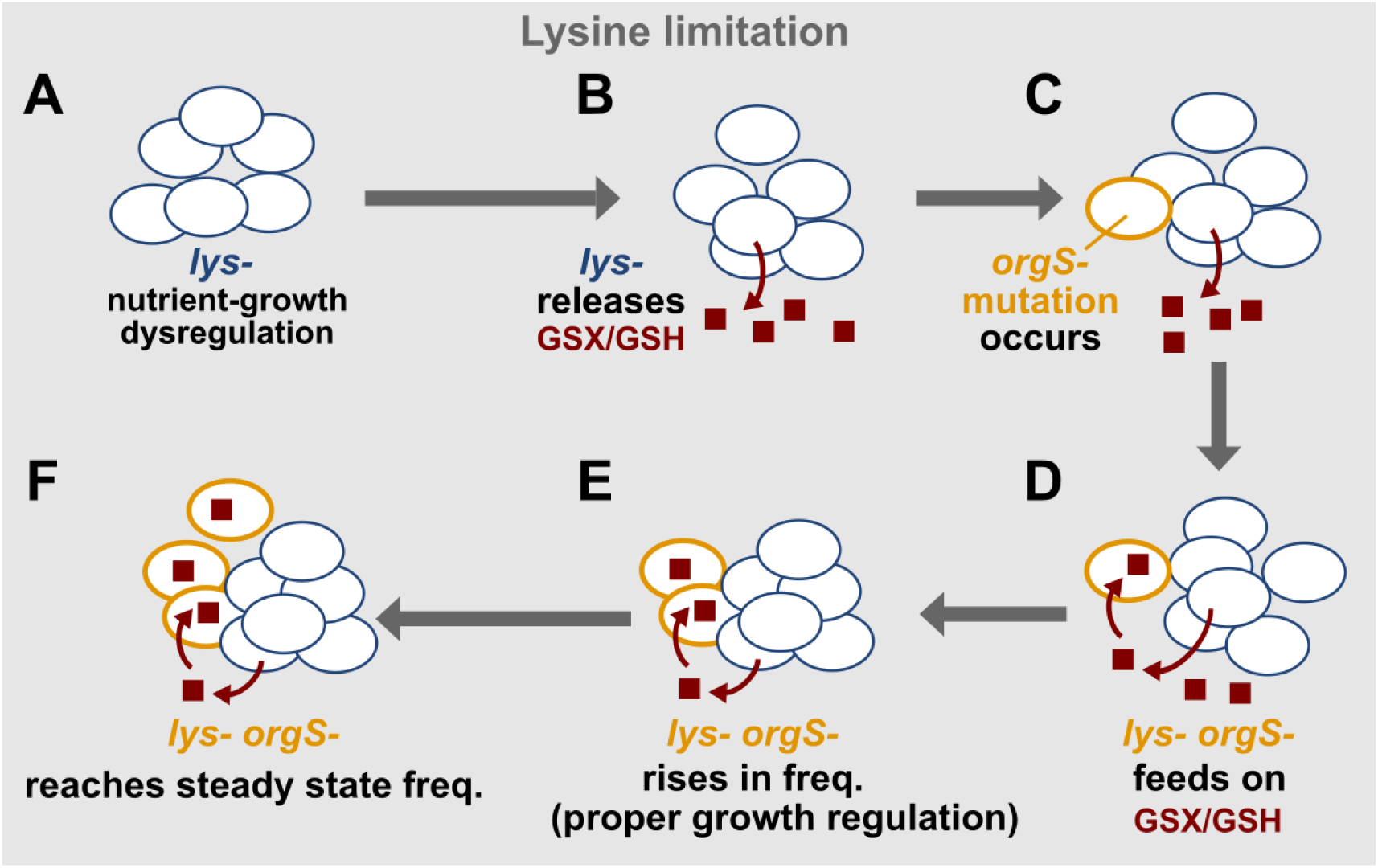
Nutrient-growth dysregulation and the evolution of an organosulfur-mediated interaction. (**A**) *lys*^*-*^ cells during lysine limitation suffered reduced autophagy and viability compared to during natural limitation (Fig 2). (**B**) These cells released organosulfurs mainly comprising GSX and, to a lesser extent, GSH (Fig 4). We hypothesize that organosulfur release is a detoxification response to nutrient-growth dysregulation, since organosulfur release was significantly reduced during glucose limitation (Fig 5B) or in excess lysine (Fig S7). (**C-D**) The released organosulfurs created a metabolic environment that could support the growth of the newly arisen *lys*^*-*^*orgS*^*-*^. (**E**) Rare *lys*^*-*^*orgS*^*-*^ mutant rose in frequency (Fig 6C) due to a fitness advantage gained by properly responding to sulfur limitation — a natural limitation. For example, compared to lysine-limited *lys*^*-*^ cells, *lys*^*-*^*orgS*^*-*^ cells doubly starved for sulfur and lysine maintained high cell viability in an autophagy-dependent fashion (Fig 6B). (**F**) Eventually, *lys*^*-*^*orgS*^*-*^ reached a steady-state frequency where organosulfur supply and consumption were balanced (Fig 6C).

The fitness advantage of *lys*^*-*^*orgS*^*-*^ over *lys*^*-*^ is presumably due to *lys*^*-*^*orgS*^*-*^ properly sensing and responding to sulfur limitation. Indeed, autophagy and low organosulfur were required for the enhanced viability of *lys*^*-*^*orgS*^*-*^ compared to *lys*^*-*^ cells during lysine starvation (Fig 6B). Consistent with this notion, mutants defective in methionine synthesis died slowly during methionine limitation (Unger and Hartwell, 1976). Furthermore, methionine auxotrophs (e.g. *met6* or *met13*) responded to methionine starvation by engaging in oxidative metabolism, similar to how cells respond to phosphate or sulfur starvation (Petti et al., 2011) (Fig 1B, red box). As *lys*^*-*^*orgS*^*-*^ cells increased in frequency, their fitness advantage over *lys*^*-*^ diminished due to competition for organosulfurs released by *lys*^*-*^ cells (Fig 6C; Fig S13). Eventually, a steady state ratio of *lys*^*-*^ *orgS*^*-*^:*lys*^*-*^ was established (Fig 6C).

The long-term dynamics of *orgS*^*-*^ is likely complex, especially in environments with fluctuating nutrient availability. Although *lys*^*-*^*orgS*^*-*^ enjoyed a frequency-dependent fitness advantage over *lys*^*-*^ during lysine limitation (Fig 6C), *lys*^*-*^*orgS*^*-*^ suffered a fitness disadvantage in excess lysine and GSH (Fig 6A). Thus, *lys*^*-*^*orgS*^*-*^ can go extinct during nutrient excess, although during lysine limitation, new *orgS*^*-*^ mutants can arise and increase in frequency to a steady state ratio. In any environment, if additional mutations acquired by *lys*^*-*^ were much more adaptive than those acquired by *lys*^*-*^*orgS*^*-*^, then *lys*^*-*^*orgS*^*-*^ can become very rare or even go extinct, similar to (Waite and Shou, 2012).

We also recovered a *gln1*^*-*^ mutation in *lys*^*-*^ cells evolving in lysine limitation (Fig 3A). Thus, lysine-limited *lys*^*-*^ cells must have released glutamine or glutamine-derivatives which supported *lys*^*-*^*gln*^*-*^ cells. Intriguingly, glutamine is the key limiting intracellular metabolite upon nitrogen limitation in *S. cerevisiae* (Boer et al., 2010). Thus, *lys*^*-*^*gln*^*-*^ might properly downregulate growth by sensing nitrogen limitation (Thevelein et al., 2000; Zaman et al., 2008), gaining a frequency-dependent fitness advantage over *lys*^*-*^ cells. Glutamine auxotroph was much rarer than organosulfur auxotrophs, possibly because glutamine consumption per cell is large with respect to glutamine release rate. A future direction would be to characterize metabolites released by cells under unnatural versus natural nutrient limitations, and to understand why metabolites are released and how released metabolites affect other cells.

### GSX and GSH release: how and why?

GSH is the major cellular redox buffer in eukaryotes. GSH carries out a variety of functions such as protecting cells against diverse forms of stress, sulfur storage and transport, and detoxification of metals and xenobiotics (Bachhawat et al., 2013). Toxic metals and xenobiotics are conjugated to glutathione in the form of GSX and subsequently exported from cells, and often GSH is co-exported (Bachhawat et al., 2013). Some stresses strongly induce genes involved in glutathione detoxification. For example, temperature shock, nitrogen depletion, and progression into stationary phase induced genes encoding glutathione precursor synthesis, glutathione S-transferase, and glutathione peroxidase (Gasch et al., 2000). Thus, glutathione detoxification is an integral part of responding to stresses including nutrient limitation.

We propose that nutrient-growth dysregulation can trigger the release of organosulfurs. Our proposition is consistent with the following observations. First, organosulfurs are barely released by *lys*^*-*^ cells grown in excess lysine (Fig S7), but rapidly released by live cells upon lysine limitation (Fig 4C, Fig 5B). In fact, the more severe the lysine limitation, the higher the organosulfur release rate (Fig S4C). Second, organosulfur release is not just a consequence of slow growth: At the same low growth rate, glucose-limited *lys*^*-*^ cells (under proper nutrient-growth regulation) released much less organosulfurs than lysine-limited *lys*^*-*^ cells (Fig 5B). Third, adenine-limited *ade*^*-*^ cells, which suffered aspects of nutrient-growth dysregulation (Fig S9) (Hart et al., 2019b), also released a significant amount of organosulfurs (Fig 5C). Together, these results suggest that nutrient-growth dysregulation is associated with, and can possibly trigger, organosulfur release.

How might organosulfurs be released? Cellular GSX and GSH are likely released by live cells rather than leaked from dead cells (Figs 4C, S6 and S8), consistent with the ATP-dependent export of GSH and GSX previously observed in yeast (Rebbeor et al., 1998). Release is presumably mediated by one or several GSX/GSH efflux pumps, including Opt1, Gex1, Gex2, and Gxa1, as well as possibly via exocytosis of vacuolar GSX/GSH (transported from cytoplasm to vacuole via Ycf1 and Bpt1) (Bachhawat et al., 2013). Yeast have been shown to upregulate glutathione transporters such as Gex1 and excrete GSH and GSX in response to cadmium-induced redox stress (Dhaoui et al., 2011). Yeast also upregulate the oligopeptide and GSX/GSH transporter Opt1 in response to different environmental stressors (Hu et al., 2017; Petti et al., 2011). We hypothesize that during nutrient-growth dysregulation, GSX conjugates were formed with cellular targets, and released for detoxification. One possibility is that cells limited for a specific metabolite upregulate the biosynthesis of that metabolite, and in an auxotroph, this upregulation may result in futile runs of the reactions preceding the missing enzyme. Since many biosynthetic pathways utilize NADPH and NADH for redox reactions, glutathione detoxification may be employed to maintain a viable redox state in the cell. A low level of GSH is co-exported with GSX, either because GSH can serve as a low-affinity substrate for GSX exporters, or because GSH export can facilitate GSX export (Bachhawat et al., 2013). Future molecular identification of the released GSX conjugates will shed light on how glutathione and nutrient-growth dysregulation might be linked.

Why might GSX and GSH be released from cells suffering nutrient-growth dysregulation? It is unlikely that cells released organosulfurs at a net cost to self. Specifically, a well-mixed environment favors self-serving changes, while a spatially structured environment can favor changes that help neighbors at a cost to self (Harcombe, 2010; Harcombe et al., 2018; Hart & Pineda et al., 2019; Momeni et al., 2013; Pande et al., 2016). This is because in a spatially structured environment, interactions are repeated and thus individuals that help neighbors to grow better will have preferential access to benefits produced by neighbors. In contrast in a well-mixed environment, individuals that contribute more are not preferentially rewarded, and thus self-serving changes are favored. Since organosulfur release occurred in a well-mixed environment, release is likely self-serving. For instance, redox buffering via GSH conjugation and the subsequent export of GSX may improve cell viability; extracellular GSH may help maintain membrane integrity (Zhang et al., 2010). To test these in the future, one could remove all genes encoding organosulfur transporters, and examine whether a mutant incapable of releasing organosulfurs is less fit than the parent strain.

### Diverse routes to the evolution of metabolic interactions

Chemical release by cells (“niche construction”) mediates diverse microbial interactions (Hillesland Kristina Linnea, 2017; Laland et al., 1999; Morris et al., 2012; Stams et al., 2006). Often, metabolites are released as a consequence of “overflow” metabolism (Basan et al., 2015; Kinnersley et al., 2009; Paczia et al., 2012; Ponomarova et al., 2017; Szenk et al., 2017). For example, yeast excretes amino acids as a consequence of nitrogen overflow, and the released amino acids in turn enable the survival of symbiotic lactic acid bacteria (Ponomarova et al., 2017). As another example, when yeast increases potassium uptake during potassium limitation, ammonium - of similar size as potassium-leaks into the cell. Since high ammonium influx is toxic, the cell detoxifies ammonium by excreting amino acids (Hess et al., 2006).

Auxotrophs are predicted to be widespread (D’Souza et al., 2014; Mee et al., 2014), and have been found in natural isolates (Beliaev et al., 2014; Carini et al., 2014; Helliwell et al., 2011; Jiang et al., 2018; Rodionova et al., 2015; Zengler and Zaramela, 2018). When nutrients are supplied by either the medium or other microbes, initially rare auxotrophic mutants can rise to a detectable frequency (D’Souza et al., 2014; D’Souza and Kost, 2016; Dykhuizen, 1978; Zamenhof and Eichhorn, 1967) via diverse mechanisms. First, at a high mutation rate (e.g. a high loss rate of plasmids that carry biosynthetic genes), auxotrophs can rise to a detectable level (Campbell et al., 2015) via mutation-selection balance: a high level of mutant is generated, and a fraction is purged by selection. Second, an auxotrophic mutation can rise to a high frequency if it happens to occur in a highly fit genetic background (“genetic hitchhiking”). Third, an auxotroph can have a fitness advantage over its non-auxotrophic (“prototrophic”) counterpart. The “energy saving” hypothesis posits that auxotrophs may gain a fitness advantage over prototrophs by saving the energy of synthesizing the essential nutrient (Zamenhof and Eichhorn, 1967). Contrary to the energy-saving hypothesis, the fitness advantage of *lys*^*-*^*orgS*^*-*^ over *lys*^*-*^ required low organosulfurs (Figs 6, S10 and S11). The fitness advantage also required autophagy (Figs 6 and S10), a hallmark of proper growth regulation during nutrient limitation. Overall, limited organosulfurs, by mimicking natural sulfur limitation and thus partially restoring property nutrient-growth regulation, allowed *lys*^*-*^*orgS*^*-*^ cells to survive lysine limitation better than *lys*^*-*^ (Fig 6).

Interestingly, nutrient-growth dysregulation has been implicated in certain cancers (Pópulo et al., 2012). Thus, understanding the evolutionary and ecological consequences of nutrient-growth imbalance in mammalian cells could be fruitful. In addition, we can view “unnatural limitation” as posing an evolutionarily novel stress that cells are ill-adapted for. We speculate that when encountering evolutionarily novel stresses (e.g. introduced by anthropogenic impacts), microbes may release metabolites that are normally not released, potentially facilitating the evolution of unexpected new ecological interactions.

## Acknowledgements

We thank Maxine Linial, Shannon Murray, Linda Breeden, and Lucas Sullivan for critically reading the manuscript, the Shou lab for discussions, Daniel Klionsky for advice on autophagy assays, and Sabrine Hedouin in Sue Biggins lab for assistance with immunoblotting.

## Methods

### Medium and strains

All strains used in this study are listed in Table S1. All yeast nomenclature follows the standard convention. For all experiments, frozen yeast strains stored at −80°C were first struck onto YPD plates (10 g/L yeast extract, 20 g/L peptone, 20 g/L glucose + 2% agar) and grown at 30°C for ∼48 hours, from which a single colony was inoculated into 3 mL of YPD and grown overnight at 30°C with agitation. All experiments were carried out within 3 days of generating the overnight culture. Minimal medium (SD) contained 6.7 g/L Difco Yeast Nitrogen Base w/o amino acids w/ ammonium sulfate and 20 g/L D-glucose, except during glucose-limitation experiments where lower levels of glucose were used as specified. Glucose starvation medium (S) comprised only 6.7 g/L Difco Yeast Nitrogen Base w/o amino acids w/ ammonium sulfate. Nitrogen starvation medium (SD-N) contained 1.7 g/L Difco Yeast Nitrogen Base without amino acids or ammonium sulfate and 20 g/L D-glucose. Depending on strain auxotrophy, SD was supplemented with lysine (164.3 µM), adenine sulfate (108.6 µM) (Guthrie and Fink, 1991), or organosulfurs (134 µM) so that cells could grow exponentially.

Strains were constructed either via yeast crosses or by homologous recombination (Guthrie and Fink, 1991; Waite and Shou, 2014). Crosses were carried out by mating parent strains, pulling diploids, sporulation, tetrad dissection and selection on suitable plates. As an example of gene deletion, the *bcy1*Δ strain (WY2527) was constructed by PCR-amplifying the KanMX resistance gene from a plasmid (WSB26 (Wach et al., 1994)) using the primers WSO671(TACAACAAGCAGATTATTTTCAAAAGACAACAGTAAGAATAAACGcagctgaagcttcgtac gc) and WSO672(GAGAAAGGAAATTCATGTGGATTTAAGATCGCTTCCCCTTTTTACataggccactagtggatct g), with a 45-basepair homology (uppercase) to the upstream and downstream region of the *BCY1* gene, respectively. The *lys*^*-*^ strain WY2490 was then transformed with the PCR product and transformants were selected on a G418 plate. Successful deletion was confirmed via a checking PCR with a primer upstream of the *BCY1* gene (WSO673 TATACTGTGCTCGGATTCCG) paired with an internal primer for the amplified KanMX cassette (WSO161 ctaaatgtacgggcgacagt).

### Evolution

Coculture evolution was described in an earlier study (Hart & Pineda et al., 2019). To revive a coculture, ∼20 µL was scooped from the frozen sample using a sterile metal spatula, diluted ∼10-fold into SD, and allowed to grow to moderate turbidity for 1-2 days. The coculture was further expanded by adding 3 mL of SD. *lys-* evolved clones were isolated by plating the coculture on rich media (YPD) agar with Hygromycin B.

For monoculture evolution, chemostat vessels (Fig S14) were placed into a core manifold with six receptacles, each with a magnetic stirrer. Reactor vessels were immobilized in receptacles by adjustable compression rings. A rubber stopper equipped with an inflow tube and a sampling needle covered the top of the vessel. Waste flowed by gravity to a waste receptacle below the device through C-Flex 0.375” ID tubing (Cole Parmer) attached to the outflow arm. Nutrient media was fed to the vessels from media reservoirs by tubing passed through a Cole Parmer MasterFlex C/L peristaltic pump controlled by a custom LabView console through a custom relay box. Media reservoirs were 1 L glass bottles capped with one-hole rubber stoppers. Tubing was fed through the stopper and allowed to hang to the bottom of the reservoir. The reservoirs were placed on Denver Instrument XP-1500 digital balances, and the actual flow rate for each vessel was determined from the rate of mass loss of the corresponding reservoir.

To create a sterile environment, initial assembly was done in autoclave trays, with vessels held in tube racks. Six reservoirs were prepared by adding 810 mL water to each bottle. Six vessels were prepared by adding a 10 mm stir bar and 20 mL growth media (SD+21 µM lysine) to each vessel. Media delivery tubing was attached between reservoirs and vessels through rubber stoppers, and waste tubing was attached to each outflow arm, with the unattached end covered by foil held in place by autoclave tape. A 1.5 mL micro-centrifuge tube was placed over the sampling needle and held in place by autoclave tape. Tubing ports were wrapped with foil as well. Each reservoir with its attached tubing was weighed, the entire assembly autoclaved, then each reservoir weighed again. Lost water was calculated and added back. Under sterile conditions, 90 mL of 10X SD and a lysine stock were added to each reservoir to reach a final lysine concentration of 21 µM. The vessels were then secured into the chemostat manifold receptacles, reservoirs placed on the scales, and tubing threaded into the pumps.

Ancestral or evolved *lys-* clones were grown in 50 mL SD + 164 µM lysine for ∼20 hours prior to inoculation. Before each experiment, growth was tracked to ensure cells were growing optimally (∼1.6 hour doubling time). When cells reached a density of ∼0.2 OD_600_, cells were washed 3 times in SD and inoculated in a chemostat vessel prefilled with SD + 21 µM lysine. After this step, chemostat pumps were turned on at a set doubling time in the custom written LabView software package. Each chemostat vessel contained ∼43 mL running volume, and was set to a target doubling time (e.g. for 7 hour doubling, flow rate is 43*ln2/7=∼4.25 mL/hr). We evolved three lines at 7-hr doubling, and three lines at 11-hr doubling. With 21 µM lysine in the reservoir, the target steady-state cell density was 7×10^6^/ mL. In reality, live cell densities varied between 4×10^6^/ml and 1.2×10^7^/ml. Periodically, 4 mL of supernatant was harvested and dispensed into a sterile 15 mL conical tube. Next, 300 µL of this cell sample was removed and kept on ice for flow cytometry analysis. The remaining 3.7 mL of supernatant was filtered through a 0.22 µm nylon filter into 500 µL aliquots and frozen at −80C. Each chemostat was sampled according to a pre-set sequence. For experiments with metabolite extraction, the chemostat vessel stopper was removed and cells from 20 mL of sample was harvested. Due to the breaking of vessel sterility, this would mark the end of the chemostat experiment.

The nutrient reservoir was refilled when necessary by injecting media through a sterile 0.2 µm filter through a 60 mL syringe. To take samples sterilely, the covering tube on the sampling needle was carefully lifted, and a sterile 5 mL syringe was attached to the needle. The needle was then wiped with 95% ethanol, and slowly pushed down so that the tip was at least ∼10 mm below the liquid level. A 5 mL sample was drawn into the syringe, the needle pulled up above the liquid surface, and an additional 1 mL of air drawn through to clear the needle of liquid residue. The syringe was then detached and the cap placed back on the needle. The samples were ejected into sterile 13 mm culture tubes for freezing and flow cytometry determination of live cell densities. In both evolution experiments, samples were frozen in 1 part 20% trehalose in 50mM sodium phosphate buffer pH 6.0 + 1 part YPD. The samples were cooled at 4°C for 15 min before frozen down at −80°C.

Whole-genome sequencing and data analysis are described in detail in (Hart & Pineda et al., 2019).

### Quantifying auxotroph frequency

Frozen cultures (2 time points from three mono-culture evolution and three co-culture evolution experiments) were revived, and clones were isolated and screened for auxotrophy. We revived frozen samples by directly plating samples on YPD (monoculture) or YPD + hygromycin (cocultures to select against the partner strain). Plates were grown at 30°C for 2∼4 days until all colonies were easily visible for picking. We observed a variety of colony sizes, and screened both large and small sized colonies when both were present. We counted large and small colonies to estimate the ratio of large:small colony-forming cells in the population, then multiplied this fraction by the fraction of auxotrophs observed in each colony size class to get a full population auxotroph frequency estimate. To screen for auxotrophy, entire colonies were inoculated into 150 µL of SD, 10 µL of which was diluted into 150 µL SD in microtiter plates and incubated overnight to deplete organosulfur carry-over or cellular organosulfur storage. In the case of some small colonies, no dilution was made as the inoculated cell density was already low enough, based on OD measurements. 10-30 µL were then diluted into a final volume of 150 µL each of SD+Lys, SD+Lys+Met, and YPD, aiming for OD∼0.005-0.03 based off an initial reading by a 96-well plate OD600 reading of the starvation plate. Plates were then incubated for 48+ hours to grow cultures to saturation, and culture turbidity (OD600) was read using a 96-well plate reader. Control wells of known *lys*^-^*orgS*^-^(WY1604) and *lys*^-^(WY2226) were included in the screening as controls. Wells that grew in SD+Lys+Met and YPD but failed to grow in SD+Lys were scored as *lys*^-^*orgS*^-^.

### Fluorescence microscopy

Fluorescence microscopy experiments and data analysis are described in detail elsewhere (Hart et al., 2019b). Briefly, the microscope is connected to a cooled CCD camera for fluorescence and transmitted light imaging. The microscope harbors a temperature-controlled chamber set to 30°C. The microscope is equipped with motorized switchable filter cubes capable of detecting a variety of fluorophores. It also has motorized stages to allow z-autofocusing and systematic xy-scanning of locations in microplate wells. Image acquisition is done with an in-house LabVIEW program, incorporating autofocusing in bright field and automatic exposure adjustment during fluorescence imaging to avoid saturation. Previous analysis (Hart et al., 2019b) has demonstrated that if fluorescence per cell is constant over time, then background-subtracted fluorescence intensity scales proportionally with live cell density, and a decrease in fluorescence intensity correlates well with cell death.

For experiments in Figs 2 and S2, WY2490 (*lys-*) and WY2527 (*lys*^*-*^*bcy1*^*-*^) were grown overnight to exponential phase in SD+164 µM lysine, washed 3 times in S medium, and starved for 3 hours at 30 °C in factory-clean 13 mm test tubes in either SD (lysine starvation) or S (lysine and carbon starvation). For imaging, ∼10,000 cells/well were inoculated into each well of a 96 well plate in the corresponding medium (rapamycin treatment was done in SD+1 µM rapamycin). The microtiter plate was imaged periodically (1∼2 hrs) under a 10x objective in a Nikon Eclipse TE-2000U inverted fluorescence microscope using an ET DsRed filter cube (Exciter: ET545/30x, Emitter: ET620/60m, Dichroic: T570LP). A similar protocol was followed for experiments in Figs 6, S9, S10 and S11, with genotypes and starvation conditions noted in the corresponding figure legends.

### Chemostats and turbidostats

Cells were grown under controlled conditions using a custom-made continuous culturing device (Fig S14A), with six channels (Fig S14C) where each can be independently operated as a chemostat or a turbidostat. When operated as a chemostat, a channel provides a limited nutrient environment where the growth rate is held constant. When operated as a turbidostat, a channel maintains a constant cell density while cells grow with an abundant supply of nutrients.

The continuous culturing device consisted of six reactor vessels (Fig S14A), each with a volume of ∼43ml determined by the height of the outflow tube (Fig S14C, E). A rubber stopper equipped with an inflow tube and a sampling needle covered the top of each vessel (Fig S14C). The vessels were placed in an aluminum mounting frame with six receptacles (Fig S14A, back), each equipped with an integrated magnetic stirrer (made from a CPU fan) and an LED-phototransistor optical detector for OD measurements. The vessels were immobilized in the receptacles by adjustable compression rings. A sampling needle passed through a short length of PharMed rubber tubing (Fig S14C). The tubing was held in place by glass tubing inserted into the stopper. Zip-ties are used to achieve the proper tightness, allowing movement of sampling needle while applying enough friction to maintain position. Waste flowed by gravity to a waste receptacle below the device through 0.375” ID tubing (Cole Parmer C-Flex) attached to the outflow arm. Nutrient media was fed to each vessel from an independent reservoir by a peristaltic pump (Welco WPM1, operated at 7V DC). The media delivery tube consisted of two sections of generic 2mm OD, 1mm ID PTFE tubing, joined by insertion into the two ends of a 17cm section of PharMed AY242409 tubing (Saint-Gobain) which was inserted into the peristaltic pump (Fig S14B). The pump was activated and deactivated by the custom LabView program through a relay box (Pencom Design, Inc. UB-RLY-ISO-EXT-LR). Depending on whether a given channel was in chemostat or turbidostat mode, the LabView program controlled pump in different ways (i.e. constant dilution rate in chemostat, or dilution to a set turbidity in turbidostat). Data for OD and flow rate data were logged in either mode of operation. In both cases, flow rate can be used to calculate growth rate. Media reservoirs (Fig S14A front) were 1 L glass bottles capped with one-hole rubber stoppers, and a section of glass tubing was used as a sleeve to prevent curling of the PTFE tubing and to keep the end of PTFE tubing touching the bottom of the reservoir. Each reservoir was placed on a digital balance (Ohaus SPX2201) with a digital interface (Ohaus Scout RS232 interface) for measurement of the volume (weight) remaining in the reservoir at any given time.

When in turbidostat mode, constant average turbidity was maintained. Specifically, the pump was activated when the measured OD was above the set point and deactivated when the OD was below set point. OD was measured using an 940nm LED (Ledtecch UT188X-81-940, driven with 50ma current) and phototransistor (Ledtech LT959X-91-0125). Each LED-phototransistor pair was tested and selected for consistent OD measurements. The LED and phototransistor were positioned by mounting holes on the aluminum metal frame, on opposite sides of the reactor vessel, 4 cm from the vessel bottom. Each phototransistor was connected to an op-amp (LM324) circuit that acted as a current to voltage converter and buffer (Fig S14B). An isolated DC-DC converter provided a regulated voltage supply for the electronics. The output voltage from the photodetector circuit was digitized using a DAQ (National Instruments USB-6009), and read by the LabView program for OD measurement. The Labview program stored the average light intensity I_0_ over the first two minutes after starting a channel as the “blank” value., The light intensity, I, was measured every ∼30s, and the OD = log_10_(I/I_0_) was calculated.

When in chemostat mode, a constant average flow rate *f* of medium into the vessel was maintained. Unlike our earlier chemostat setup (Skelding et al., 2018), here constant flow rate was achieved via a scale (Ohaus SPX2201) which constantly weighed its associated reservoir, and the reading was acquired through an RS232 interface (Ohaus Scout RS232). The initial scale reading, *M*_*initial*_, was recorded when the chemostat channel was activated or reset. This was used with the current scale reading *M*_*current*_ to calculate the total volume pumped from the reservoir to the vessel as 1*ml/g * (M*_*current*_ *-M*_*initial*_). The target volume that should have flown from the reservoir to the vessel at the current time was calculated according to the pre-set flow rate. If the total volume was less than the target, the pump was activated, and otherwise, the pump was deactivated. This provided the correct average flow rate. The flow rate was chosen for the desired doubling time, *f = ln(2) * V / T*_*D*_, where *V* is the volume of the vessel and *T*_*D*_ the doubling time. Vessel volumes were measured by weighing empty vessel, and then weighed again when filled to the spillover point (Fig S14E), giving an average value of 43ml. Individual flow rates were determined using individual vessel volumes. Volume measurements are limited by the minimum waste tube drop size of ∼0.5 ml, which is constrained by surface tension. Scale readings were logged, providing a measure of flow rate (Fig S14D).

### Bioassays

Most bioassays are yield-based. Various tester strains were used in this study (specified in each experiment; Table S1), but preparation for all yield-based bioassays was the same. Strains were grown for ∼16 hours in SD and any required supplements. During this time, growth rate was tracked to ensure cells were doubling as expected (1.6 to 3 hour doubling depending on the strain/condition). After this time, cells were washed 3 times with 3 mL SD and starved for at least 3 hours at 30°C in 3 mL SD in a factory-clean 13 mm test tube (to prevent inadvertent nutrient contamination). This was done to deplete cells of vacuolar stores of nutrients. Finally, ∼1000 cells/well were inoculated into a final volume of 150 µL into a flat-bottomed 96 well plate of either metabolite standards or supernatants, supplemented with SD+1x lysine. For each auxotrophic strain, SD+1x lysine supplemented with various known concentrations glutathione were used to establish a standard curve that related organosulfur concentration (in terms of fmole GSH equivalent) to final turbidity (Fig S5). Turbidity achieved in a supernatant was then used to infer organosulfur concentration in the supernatant. Plates were wrapped with parafilm to prevent evaporation and incubated at 30°C for 2-3 days. We re-suspended cells using a Thermo Scientific Teleshake (setting #5 for ∼1 min) and read culture turbidity using a BioTek Synergy MX plate reader. In a rate-based bioassay (Fig S4B), mCherry tagged yeast strain auxotrophic for organosulfur (WY2035) was pre-grown in SD +164 µM Lys +134 µM Met and growth rate was tracked by optical density to ensure the cell was growing as expected. Next, cells were washed 3 times in SD lacking supplements, and starved for at least 3 hours in factory-clean 13 mm test tubes. OD was measured again and cells were inoculated to roughly 1000 cells /well in a 96 well plate in a total volume of 300 µL. The well was filled with either known quantities of organosulfur (methionine or glutathione) or harvested supernatants, both supplemented into SD + 164 µM lysine. The 96 well plate was measured in the same manner as previously outlined in the “Fluorescence microscopy” section.

Maximal grow rate was calculated by measuring the slope of ln(Normalized Intensity) against time. For each sliding window of 4 time points, slope is calculated and if it exceeds the current max slope for the well, it is chosen as the new maximum. To ensure that no estimation occurs when other metabolites, such as glucose, could be limiting, we restricted analysis to data at 25% maximal intensity to ensure that cells had at least 2 doublings beyond when they are theoretically growing maximally. For this study, maximal growth rates were used to estimate approximate niche size, the logic being that cells should grow faster if there is a larger organosulfur niche.

### Flow cytometry

Detailed description can be found elsewhere (Hart et al., 2019a). Population compositions were measured by flow cytometry using a DxP10 (Cytek). Fluorescent beads of known concentration (as determined by hemocytometer) were added to determine cell densities. A final 1:20,000 dilution of ToPro3 (Molecular Probes T-3605) was used for each sample to determine live and dead cell densities. Analysis using FlowJo software showed obvious clustering of live and dead cells in the ToPro3 RedFL1 channel, with dead cells having a RedFL1 signal of >10^3^. Dead cell densities typically were never higher than 10% in all conditions tested.

### Metabolite extraction

Metabolite extraction for intracellular organosulfur quantification was adapted from (Rosebrock and Caudy, 2017). Briefly, 20 mL of chemostat populations were harvested with disposable 25 mL pipettes, and rapidly vacuum filtered onto precut 0.45 µm Magna nylon filters (GVS Life Sciences, USA). Using ethanol-cleaned forceps, the filter was then quickly submerged into 3 mL ice-cold extraction mixture— 40% (v/v) acetonitrile, 40% (v/v) methanol, and 20% (v/v) distilled water—held in a sterile 5 mL centrifuge tube. All reagents were HPLC grade and all extraction buffer mixtures were made fresh before each extraction. The centrifuge tube was capped and quickly vortexed to remove all cells. The entire process took less than 25 seconds, with the time between populations being filtered and submerged in extraction buffer being less than 10 seconds. After all populations had been harvested, extracts were frozen at −80 °C until solid, transferred to ice and allowed to thaw. After the samples had thawed, they were incubated on ice for 10 minutes and vortexed once every ∼3 minutes and returned to −80°C for refreezing (a single ‘freeze-thaw’ cycle). After 3 freeze-thaw cycles, 1.5 mL of sample was harvested and transferred to a new 1.5 mL micro-centrifuge tube, and centrifuged at 13,0000 rpm for 2 minutes at 4°C to pellet the cell debris. The extract was removed and the remaining cell pellet was extracted again with 1.5 mL of extraction mixture, and spun down. The final result was 3 mL of extracted metabolites that was stored at −80°C and analyzed by HPLC less than 48 hours after extraction. To check that a majority of metabolites were extracted, 100 µL of fresh extraction buffer was added to the collected cell debris, vortexed vigorously, and collected by centrifugation. This 100 µL ‘second extract’ was also analyzed for glutathione by HPLC. On average, the amount of glutathione in the second extract was < 2% of the amount extracted initially in the 3 mL extraction.

For the extracts used in bioassays, a similar protocol was followed except 0.45 µm nitrocellulose membranes (Bio-Rad) were used and the extraction mixture was dried off using the low temperature setting on a speed-vac and the dehydrated components were resuspended in sterile distilled water. From the total amount of metabolites in the sample and the total number of cells used to extract metabolites, we can calculate the average amount of metabolite per cell.

### Analytical chemistry quantification of glutathione (GSH) and glutathione S-conjugates (GSX)

#### HPLC

Reduced glutathione was derivatized using a thiol-specific probe first described by (Yi et al., 2009), called Thiol Probe IV (EMD Millipore) to make a fluorescent glutathione conjugate. The compound reacts readily with free-thiols, though at different rates. For quantifying glutathione, 270 µL of sample or GSH standard in SD was added to 30 µL of 833 mM HEPES buffer, pH 7.8. This was done to raise the pH of the sample to a basic level, which facilitates the reaction. Next, the probe (dissolved in DMSO and stored in 50 µL aliquots at −20 C), was added to a final concentration of 100 µM, which is in excess of glutathione by at least 10-fold. The reaction was performed at room temperature in the dark (the probe is light-sensitive) in a 96-well plate for 20 minutes. After this, 8.4 µL of 2M HCl was added to rapidly quench the reaction by lowering the pH to ∼2. This also stabilizes the fluorescent conjugate. The entire sample was then added to a 250 µL small volume pulled point class insert (Agilent Part No: 5183-2085) to facilitate autosampler needle access. The small volume insert with sample was then placed inside a dark-brown 1.5 mL autosampler vial (Shimadzu Part No: 228-45450-91) and capped with a fresh 9mm screw cap with PTFE septum (Shimadzu Part No: 228-45454-91).

Derivatized glutathione was separated and identified using reverse phase chromatography. 10 µL of the reaction mixture was injected onto Synergi 4 µM Hydro-RP 80 Å LC Column 150 × 4.6 mm (Phenomenex, Part No: 00F-4375-E0) fitted with a SecurityGuard Cartridges AQ C18 4 × 3.00 mm ID (Phenomenex, Part No: AJO-7511) in a SecuirtyGuard Cartridge Holder (Phenomenex, Part No: KJ0-4282). The SecurityGuard (pre-column) was periodically replaced whenever pressure reading exceeded the manufacturer’s specifications. Glutathione was eluted from the column with a mobile phase gradient of filtered Millipore water (Solution A) and acetonitrile (Solution B, HPLC grade). The Millipore water was filtered through a 0.22 µM filter prior to use. Additionally, before each run the column was equilibrated for 30 minutes with 1% Solution B. The % solution B followed the following program for each injection: 0 min 1%, 10 min 14%, 10.01 min 1%, and 15 min 1%, corresponding to a gradual increase to 14% Solution B over 10 minutes, followed by a re-equilibration with 1% Solution B. The column was maintained at a running temperature of 25 °C in a Nexera X2 CTO-20A oven. Flow rate was 1 mL /min. Under these conditions, glutathione eluted at ∼ 7 minutes, with slight run-to-run variation. Fluorescent glutathione was detected by excitation at 400 nm and emission at 465 nm. After each run, the column was washed and stored per manufacturer’s instructions.

Analysis of HPLC data was done using the R Statistical Language with custom written software for peak-picking, baseline correction, plotting, and area estimation, which is freely available at https://bitbucket.org/robingreen525/hplc_rscripts/src/master/. Raw data for each sample run was exported to a text file and parsed in the RStudio environment. Emission data (at 465 nm) was culled to restrict analysis from 6.5 to 8 minutes. Next, the script identified a local maximal peak, which for concentrations above 0.01 µM glutathione always corresponding to glutathione. Anything lower was indistinguishable from the background. Next, the script identified local minima on both sides of the glutathione peak and drew a baseline that connected the two. The formula (y=mx+b) for this line was calculated and the baseline was ‘corrected’ by subtracting the emission spectrum value for each point against the y value of the calculated formula for the same point. To quantify the concentration of glutathione in a sample, known concentrations of GSH in SD were subjected to the above procedure. A standard curve of 0.03 to 1 µM was typically used and showed little variability between experiments. A linear regression model of peak area against concentration of reduced glutathione was built using the lm function of the stats package. Samples with areas within the dynamic range (0.03-1 µM GSH) were back-calculated using the linear regression model. Comparing HPLC traces of the same derivatized sample over 24 hours shows that glutathione peak area is within 10% of all replicates.

#### LC-MS

Supernatants were shipped over-night on dry ice to the Rabinowitz lab at Princeton University. Stable isotope compound [2-^13^C, ^15^N] Glutathione (GSH) was obtained from Cambridge Isotope Laboratories. HPLC-grade water, methanol, and acetonitrile were obtained from ThermoFisher Scientific. Supernatant sample was thawed at room temperature and 30 µL of the supernatant together with 5 µL of 10 µM 2-^13^C+^15^N labeled GSH was transferred into a 1.5 mL centrifuge tube. The samples were either run directly to measure GSH only, or first treated with TCEP to reduce GSX to GSH and then measure the total GSH. For those samples with TCEP treatment, 5 µL of 60 g/L tris(2-carboxyethyl)phosphine solution (TCEP, reducing reagent) was added into the sample. The resulting mixture was vortexed and incubated for 20 min at room temperature. Afterward, 10 µL of 15% NH_4_HCO_3_ (w/v) was introduced to neutralize the pH of the solvent. The solution was dried down under N_2_ flow and resuspended in 50 µL 40:40:20 (methanol/acetonitrile/water) solvent and kept at 4 **°**C in an autosampler.

Samples were analyzed using a Q Exactive Plus mass spectrometer coupled to Vanquish UHPLC system (Thermo Scientific). LC separation was achieved using a XBridge BEH Amide column (2.1 mm x 150 mm, 2.5 µm particle size, 130 Å pore size; Waters, Milford, MA) using a gradient of solvent A (20 mM ammonium acetate + 20mM ammonium hydroxide in 95:5 water: acetonitrile, pH 9.45) and solvent B (acetonitrile). Flow rate was 150 µl/min. The gradient was: 0 min, 90% B; 2 min, 90% B; 5 min, 50% B; 10 min, 0% B; 13.5 min, 0% B; 15 min, 90% B; 20 min, 90% B. Column temperature is 25°C and injection volume is 10 µL. Mass spectrometer parameters are: positive ion mode, resolution 140,000 at m/z 200, scan range m/z 290-650, AGC target 3E6, Maximum injection time 200 ms. Quantitation of Glutathione concentrations in samples were achieved by comparing the peak areas of glutathione to those of ^13^C-GSH. Data were analyzed using the MAVEN software (Melamud et al., 2010).

### Competition

To quantify multiple competition replicates at multiple initial strain ratios, we used the coculture system to mimic the lysine-limited environment, especially since similar mutations in coculture and monoculture lysine-limited chemostats meant that the environments were similar. To do so, WY1340 (the purine requiring/lysine releasing strain in the RM11 background) was grown to exponential phase overnight in SD+134 µM adenine, washed 3 times with SD to remove adenine, and starved for 24 hours to deplete vacuolar storage. During this starvation, WY2072/2073 (BFP *met10* evolved clones) and WY2429 (mCherry *MET10* evolved clone) was grown overnight in SD + 134 µM methionine to exponential phase, washed 3 times, washed 3 times with SD to remove excess methionine and lysine. Next, WY2072/3 and WY2429 were mixed in ratios of 1:100, 1:10, 1:1, and 10:1 to a final OD_600_ of 0.1. This mixture of populations was then added 1:1 with WY1340 to a final OD_600_ of 0.03. This was considered generation 0. Populations were monitored for growth by measuring optical density over time and periodically diluted back to OD600 0.03 (OD_600_ was never greater than 0.45 to ensure no additional metabolites from SD was limiting). The OD_600_ data was used to back calculate total generations in the experiment. Periodically, 100 µL of the culture was sampled for flow cytometry to track strain ratios. Experiments were performed until the strain ratio stabilized.

### Autophagy Assay

Autophagy activities were measured using the GFP-Atg8 cleavage assay (Xie et al., 2008). Yeast strains with *ura3* deletion in *lys*^*-*^ and *lys*^*-*^*met17*^*-*^ background were generated via crosses and transformed with GFP-Atg8 plasmid (Addgene 49425) to generate the two strains used in the autophagy assays—WY2520 (*lys*^*-*^) and WY2521 (*lys*^*-*^*orgS*^*-*^). This plasmid expresses ATG8 with an N-terminal GFP tag under the endogenous promoter in a pRS416 vector with a *URA3* selection markers (Guan et al Mol Biol Cell. 2001 Dec;12(12):3821-38). For every experiment, WY2520 and WY2521 were streaked on SC-Ura (Guthrie and Fink, 1991) plates from frozen stocks and saturated overnights were grown from single colonies in SC-Ura medium. Cultures of 25-50 mL volume were inoculated from the overnights in SD + Lysine (WY2520) or SD + Lysine + GSH (WY2521) in conical flasks and grown for 18-20 hours at 30°C to a desirable cell density. In accordance with published protocols, the initial trials aimed at a starting OD of 0.7-1.0 for starvation. However, we observed that high cell densities could result in higher GFP-Atg8 cleavage in unstarved cells. Thus, in subsequent trials, starvation was initiated at an OD in the range of 0.2-0.6. The cells were pelleted in 50 mL falcon tubes and washed thrice in sterile milliQ water. After the washes, cells were resuspended in the starvation medium of choice (see details below) in factory-clean tubes at an OD of 0.1-0.2. For the conditions where cells did not arrest growth upon starvation (primarily organosulfur starvation), cultures were periodically diluted to minimize the influence of secondary nutrient limitations caused by high cell densities. For time-course analysis, starvation was carried out in 10 mL cultures in 18 mm tubes and 2 mL of the culture was withdrawn for analysis at 24, 48 and 72 hours from the initiation of starvation. For assays with a single time-point sampling, starvation was carried out in 3 mL cultures in 13 mm tubes. *lys*^*-*^ cells were starved for 4 or 8 hours with comparable results observed for both treatments. *lys*^*-*^*orgS*^*-*^ cells were starved for 72 hours after the time-course analysis revealed that the influence of organosulfur starvation is only discernable after 48 hours in the GFP-Atg8 cleavage assay.

Starvation media for different treatments are described here. For *lys*^*-*^ cells, lysine starvation was carried out in SD medium; nitrogen starvations were carried out in SD-N medium either supplemented with 164 µM lysine (only nitrogen starvation) or lacking it (double starvation for lysine and nitrogen). For *lys*^*-*^*orgS*^*-*^ cells, double starvation for lysine and organosulfur was carried out in SD medium, lysine starvation was carried out in SD + 134 µM GSH and organosulfur starvation was carried out in SD + 164 µM lysine.

Sample preparation was carried out as suggested in (Huang et al., 2014) with minor modifications. Cells from 2-4 ml of culture were pelleted in a microcentrifuge tube at 5,000 g for 3 minutes. Cell pellets were flash frozen in liquid nitrogen and stored in −20°C till all samples had been collected for an experiment. For cell lysis, pellets were resuspended in 1 mL of ice-cold 10% Trichloroacetic acid (TCA) and allowed to stand in a cold metal block on ice for 30-40 minutes. Proteins were pelleted at 16,000 g for 1 minute at 4°C. The pellets were resuspended in 1 mL cold Acetone by vortexing and bath sonication and pelleted again by centrifugation. The acetone-wash was repeated once and pellets were allowed to air-dry for 5 minutes before resuspending in SDS-PAGE sample buffer (0.1 M Tris-HCl pH 7.5, 2 % w/v SDS, 10 % v/v glycerol, 20 mM DTT). Based on the OD measured at the time of sample collection, the sample buffer volume was adjusted to attain comparable cells/µL in each sample. Acid-washed glass beads (425-600 µm; Sigma G8771) were added to each tube, roughly equivalent to half the sample volume, and the pellet was resuspended by bead beating for 35 seconds. After centrifugation for 20 minutes to allow the foam to settle, the samples were heated in a 95°C metal block for 10 minutes. After a 3-minute centrifugation at 5,000 g, samples were run on a 12.5% acrylamide gel for 50 minutes at a constant current of 30 mA. The bands were transferred onto a 0.2 µm PVDF membrane (Bio-Rad Trans-blot 162-0184) using a wet transfer protocol at a constant voltage of 60 V for 2 hours in the cold room. For immunoblotting, the membrane was incubated overnight at 4°C with an anti-GFP monoclonal primary (JL-8; Clontech 632381) at 1:5,000 dilution, followed by a 45-minute room temperature incubation with a horseradish peroxidase-conjugated anti-mouse secondary (GE Biosciences) at 1:10,000 dilution. Antibodies were detected using the SuperSignal West Dura Chemiluminescent Substrate (Thermo Scientific). The pre-installed Gel Analyzer plugin on ImageJ was used for quantification of bands.

## Figures

**Fig S1.**
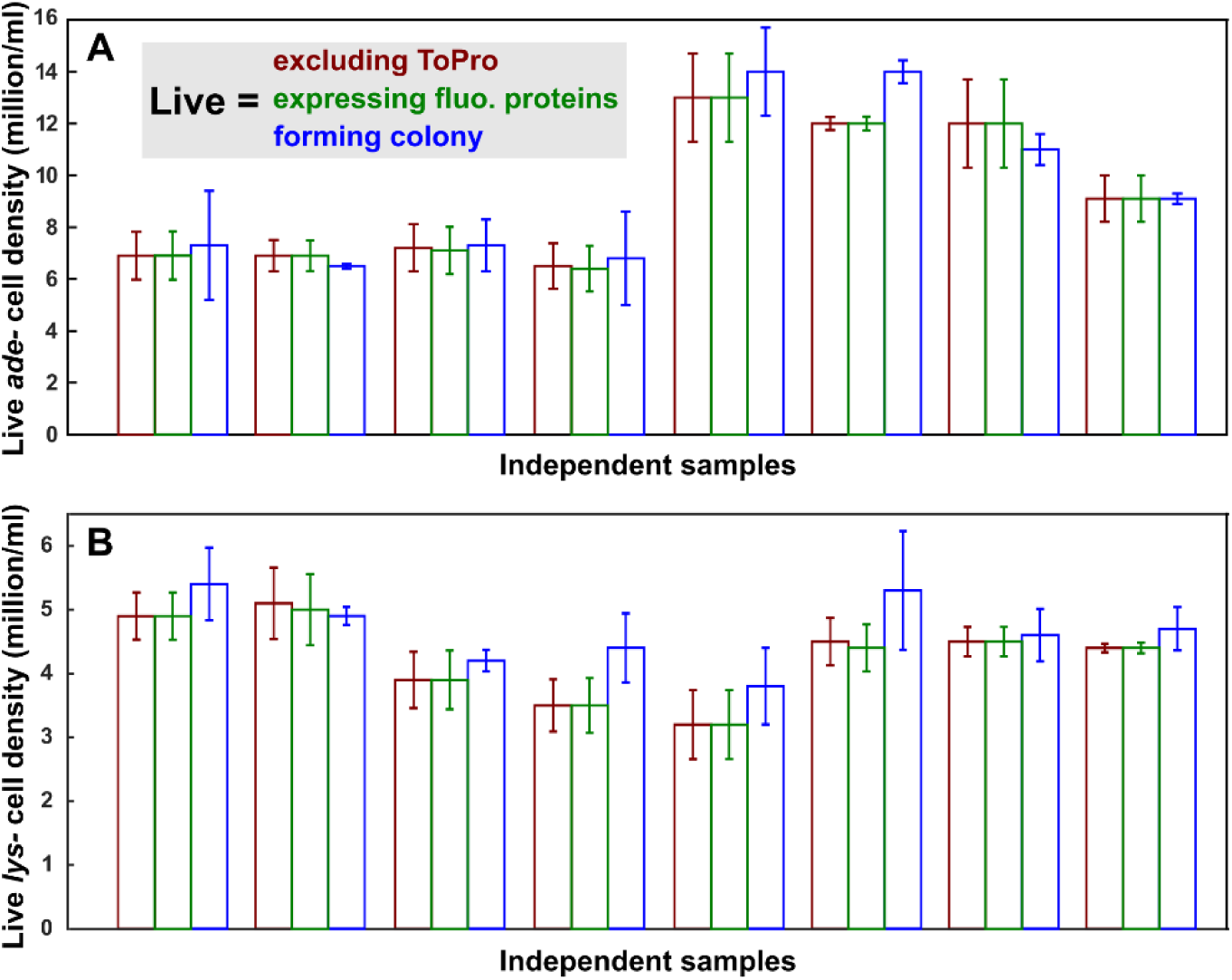
Decrease in total fluorescence can be used as a proxy for loss of cell viability. Loss of fluorescence represents cell death, as supported by three lines of evidence. First, in time lapse movies, as a cell ruptured, its fluorescence abruptly disappeared (Supplementary Movies 3 and 4 in Hart et al., 2019b). Second, regardless of whether live cells were defined as fluorescent protein-expressing cells, colony-forming cells, or cells with intact membrane and thus capable of excluding the nucleic acid dye ToPro3, we obtained similar quantification of live cell densities. Specifically, adenine auxotrophic (**A**) and lysine auxotrophic (**B**) cells expressing fluorescent proteins were grown in adenine-limiting and lysine-limiting chemostats, respectively (Hart et al., 2019a; Skelding et al., 2018). After cell density had reached a steady state, a sample was taken where a portion was plated on rich medium, and live cell density was quantified from colony counts. Another portion of the sample was analyzed by flow cytometry after being stained with ToPro3, a nucleic acid dye that can only enter cells with compromised membrane integrity. Live cell density was then quantified either from cells that excluded ToPro or from cells that expressed fluorescent proteins. Each comparison was for an independently run chemostat. We can see that the three assays generated similar quantifications of live cell densities. Third, death rate quantified from the decline in total fluorescence during nutrient starvation was similar to that quantified from the decline in ToPro3-negative live cells (Hart et al., 2019b).

**Fig S2.**
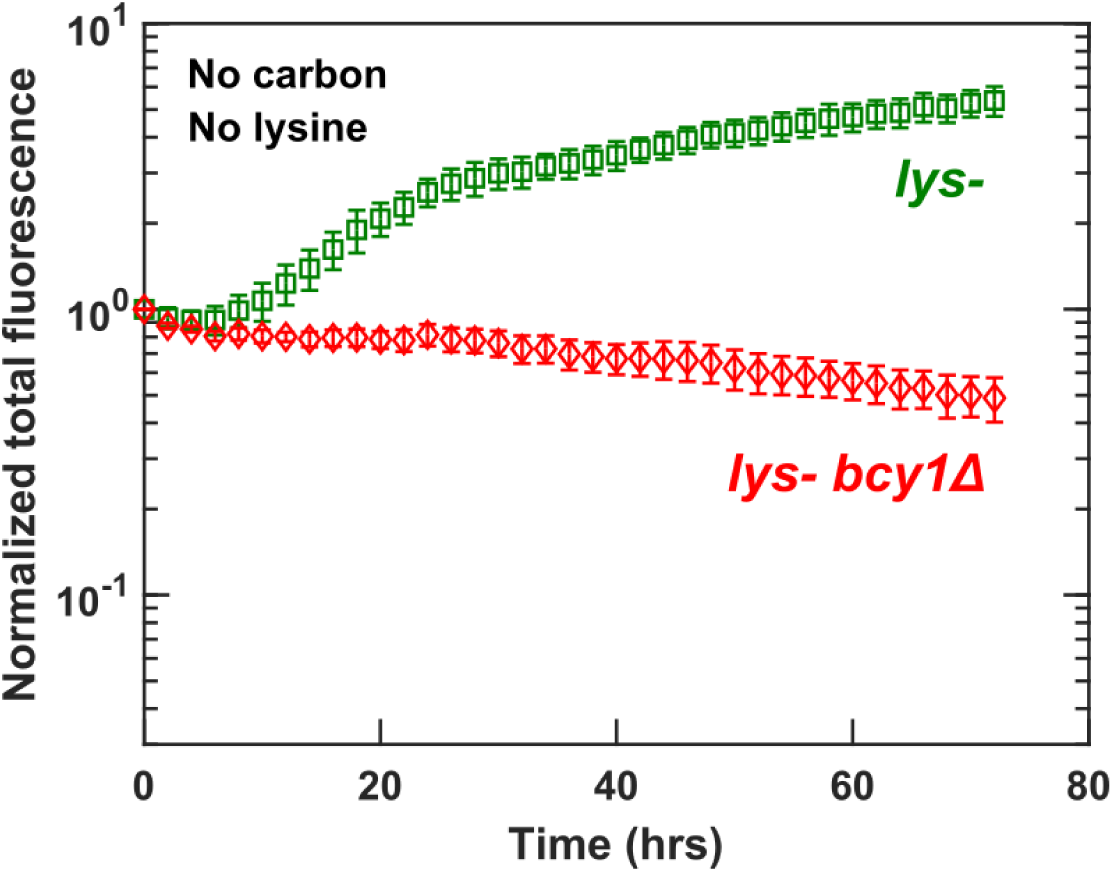
Cell viability during nutrient starvation is reduced by activating growth. Bcy1 inhibits the Ras/PKA growth-activating pathway, and thus *bcy1*Δ suffers overactive growth. Compared to *lys*^*-*^ cells (WY2490) under dual starvation for lysine and glucose, *lys*^*-*^*bcy1*Δ (WY2527) cells suffered reduced cell viability. Exponentially growing cells were washed and starved for glucose and lysine for 3 hrs, and cultured and imaged in minimal medium without glucose or lysine (Methods “Fluorescence microscopy”). The initial increase in the fluorescence of *lys*^*-*^ cells was due to cells becoming brighter. Error bars correspond to two standard deviations for 6 replicate wells.

**Fig S3.**
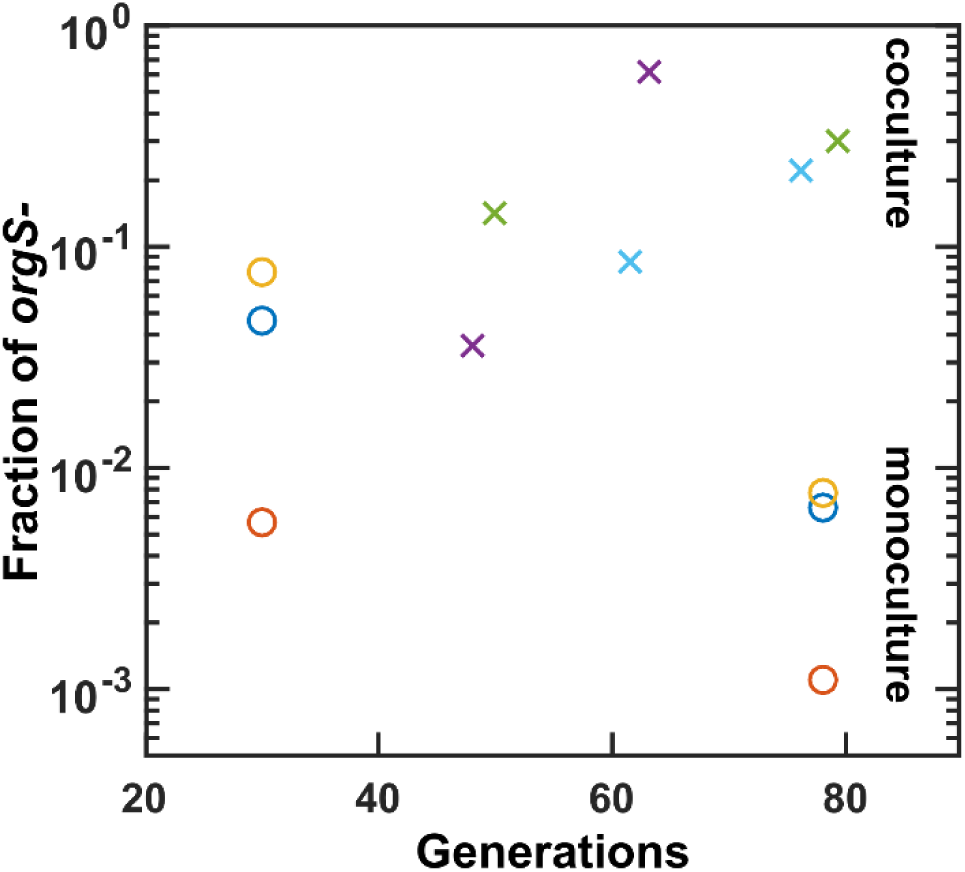
*lys*^-^*orgS*^-^repeatedly rose to detectable frequency when *lys-* cells were limited for lysine. *lys-* cells (WY1335) were either cocultured (crosses) with a lysine-releasing strain (WY1340) or cultured alone in lysine-limited chemostats (circles). Cultures were frozen periodically in 1:1 volume of YPD: 20% trehalose in 50mM NaH_2_PO_4_ (pH 6.0). We revived frozen samples by directly plating samples on YPD or, in case of cocultures, YPD + hygromycin to select against the partner strain. We observed a variety of colony sizes, and since large and small colonies had different percentages of *lys*^-^*orgS*^-^, we quantified both types and calculated the overall *orgS-* in the population (Methods “Quantifying auxotroph frequency”). Colonies that grew in YPD and SD+Lys+Met but failed to grow in SD+Lys were scored as *lys*^-^*orgS*^-^. Different colors indicate independent evolution lines. The fraction of *lys*^-^*orgS*^-^is much lower in monocultures compared to cocultures. One explanation is that in coculture experiments, the lysine-releasing *ade*^*-*^ partner strain also released organosulfurs (Fig 5C).

**Fig S4.**
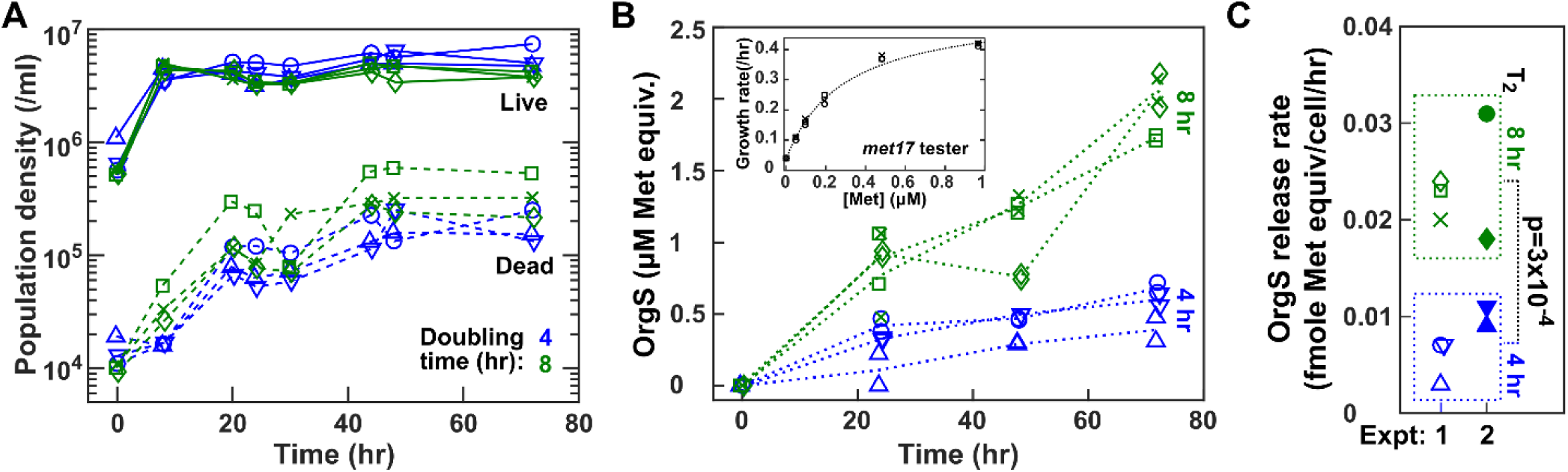
Rapid accumulation of organosulfurs upon lysine limitation. Ancestral *lys-* cells (WY1335) were grown to exponential phase in SD supplemented with excess lysine, then washed free of lysine, and inoculated into replicate lysine-limited chemostats (different symbols) with 8-hr doubling time (green) or 4-hr doubling time (blue). Periodically, culture supernatants were quickly sampled, filtered, and frozen at −80°C to preserve the redox states of released compounds. (**A**) Population dynamics of live (fluorescent) and dead (non-fluorescent) cells in chemostats, as quantified by flow cytometry (Methods, “Flow cytometry”). (**B**) Organosulfurs in supernatants were measured as “methionine equivalents” by comparing growth rates of a *met17*^*-*^ tester strain (WY2035) in supernatants fortified by SD versus in SD supplemented with known concentrations of methionine (standard curve in inset). Since the growth rate of tester cells can be affected by factors other than organosulfurs (e.g. pH), the organosulfur niche estimated by a rate-based bioassay can differ from that estimated from a turbidity-based bioassay (e.g. Fig 5B). Nevertheless, this assay motivated the LC-MS experiments that identified GSH as one of the released organosulfurs (Fig 4A). (**C**) The release rate of organosulfurs is higher in 8-hr doubling chemostats compared to 4-hr doubling chemostats. Release rates were calculated from the first 24 hrs from independent chemostats run in two experiments. If organosulfurs are the major factor affecting *met17* growth rate, then the organosulfur release rate by *lys*^*-*^ cells in 8-hr chemostats are significantly higher than that in 4-hr chemostats by t-test.

**Fig S5.**
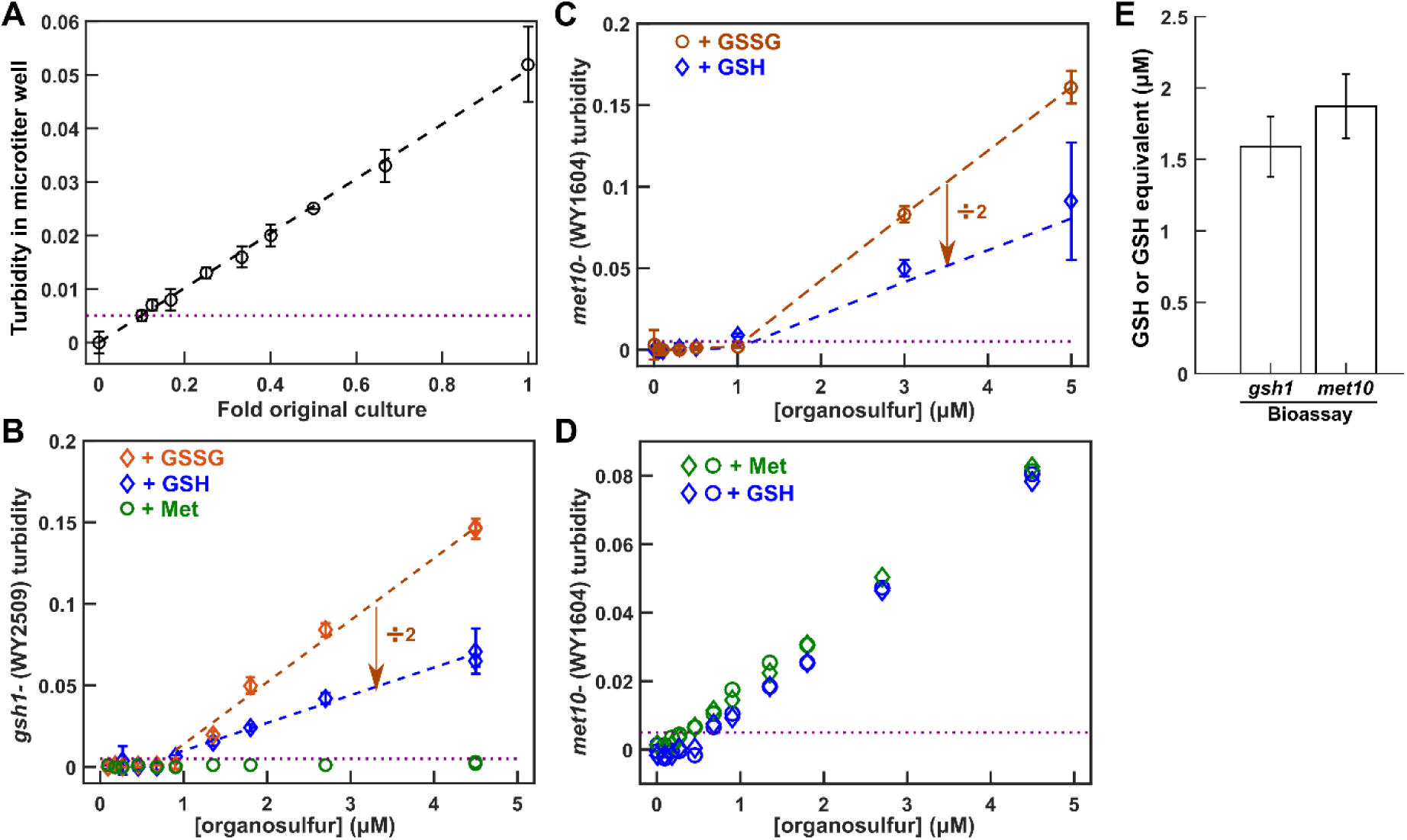
Turbidity-based yield bioassays of organosulfurs. (**A**) The sensitivity of turbidity (OD_600_) measurements in a micro-titer plate. Purple dotted line marks the lower-bound of turbidity reading that we accepted as valid data, and is plotted in **B-D**. (**B**) The turbidity of a *gsh1*^*-*^ strain (WY2509) increases linearly with [GSH] and [GSSG] within a range, but does not increase when supplemented with methionine. Since reduction of one GSSG molecule generates two GSH molecules, culture maximal turbidity in GSSG should be twice as much as in equal-molar GSH. This is indeed observed. (**C, D**) *met10*^-^ can use methionine, GSH, and oxidized glutathione (GSSG). (**E**) *gsh1*^*-*^ and *met10*^*-*^ grew to similar levels in supernatants of *lys*^*-*^ cells grown in lysine-limited chemostats.

**Fig S6.**
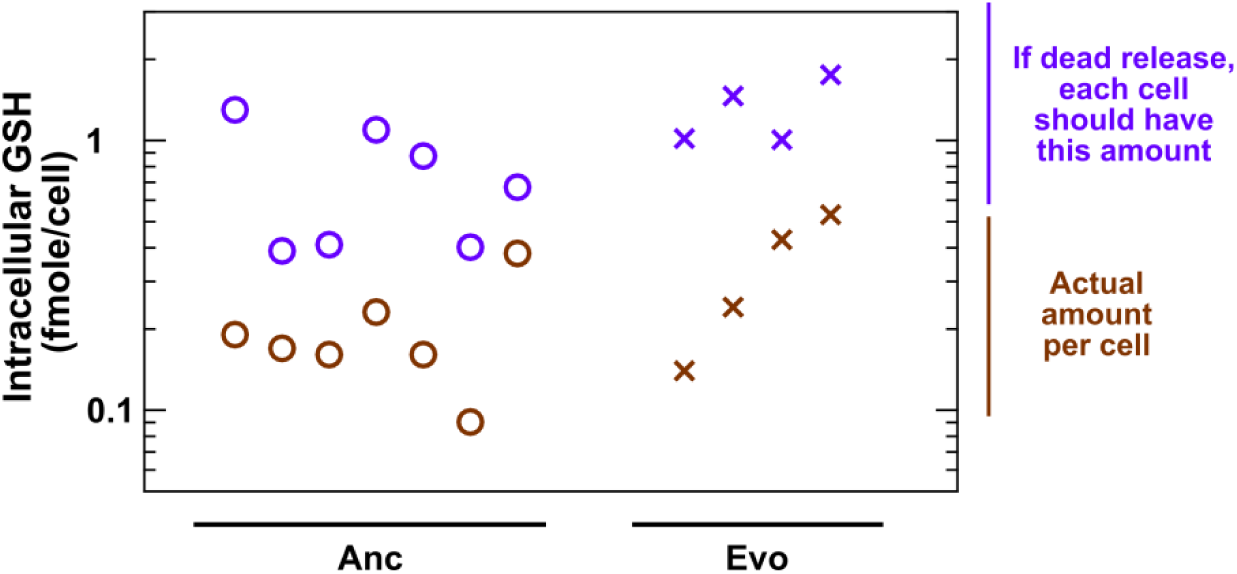
GSH is likely released by live ancestral and evolved *lys2*^*-*^ cells. Ancestral (circles; WY1335) and evolved (crosses; WY2429) *lys*^*-*^ cells were cultured in lysine-limited chemostats (doubling time 8 hrs). Evolved cells contain an *ecm21* mutation and Chromosome 14 duplication, and thus exhibit improved affinity for lysine. Intracellular metabolites were extracted from cells to quantify fmole GSH/cell (brown). We quantified dead cell density and the concentrations of GSH in culture supernatants. We then calculated the theoretical amount that would need to be inside an average cell for cell lysis alone to explain the supernatant concentrations (purple). Since the theoretical amount was higher than the actual amount in all experiments, GSH is likely released by live cells. GSH was quantified using HPLC, and dead cell density was quantified using flow cytometry (Methods). Here, each column corresponds to an experiment. Day-to-day variations exist, but the trend is clear across days.

**Fig S7.**
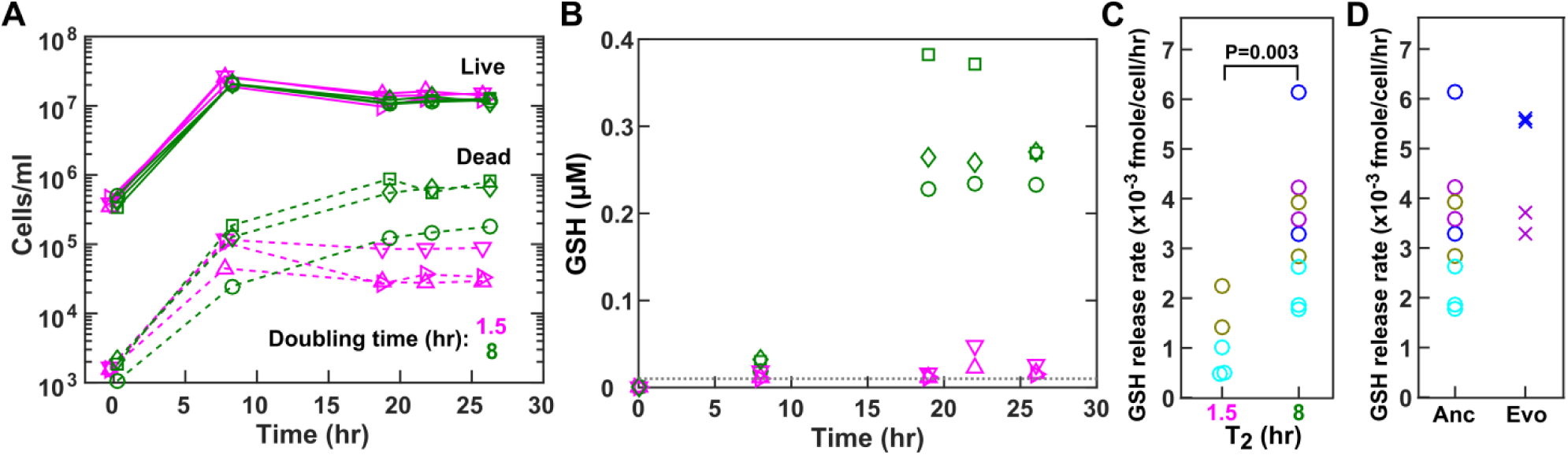
Lysine limitation increases GSH release rate. (**A-C**) Ancestral *lys2*^*-*^ cells (WY1335) were cultured in excess lysine (turbidostats at 1.5-hr doubling time; magenta) or limited lysine (lysine-limited chemostats at 8-hr doubling time; green). (**A**) Live and dead population densities were quantified using flow cytometry (Methods, “Flow cytometry”), and (**B**) supernatant GSH concentrations were quantified using a fluorescence-based HPLC assay (Yi et al., 2009) (Methods “HPLC”). Different symbols represent independent experiments. The dotted line in **B** marks the sensitivity of the HPLC assay. (**C**) Higher GSH release rate during lysine limitation. To calculate the release rate of GSH per live cell in chemostats, we used a previously described method (Hart et al., 2019a). Specifically, the steady state concentration of glutathione in a chemostat (B) was divided by the live cell density (A) and multiplied by dilution rate (/hr). P-value was derived from a one-tailed t-test assuming equal variance. Here, we used HPLC to assay GSH instead of bioassay for total organosulfurs, because the latter assay was much less sensitive. (**D**) GSH release rates were comparable between the ancestral *lys2*^*-*^ (circles) and an evolved *lys2*^*-*^ clone (crosses; WY2429 which contains an *ecm21* mutation and Chromosome 14 duplication and thus exhibits improved affinity for lysine). Here, cells were grown in lysine-limited chemostats (8-hr doubling). In (**C**) and (**D**), different colors correspond to experiments done on different days, and each symbol represents an independent culture.

**Fig S8.**
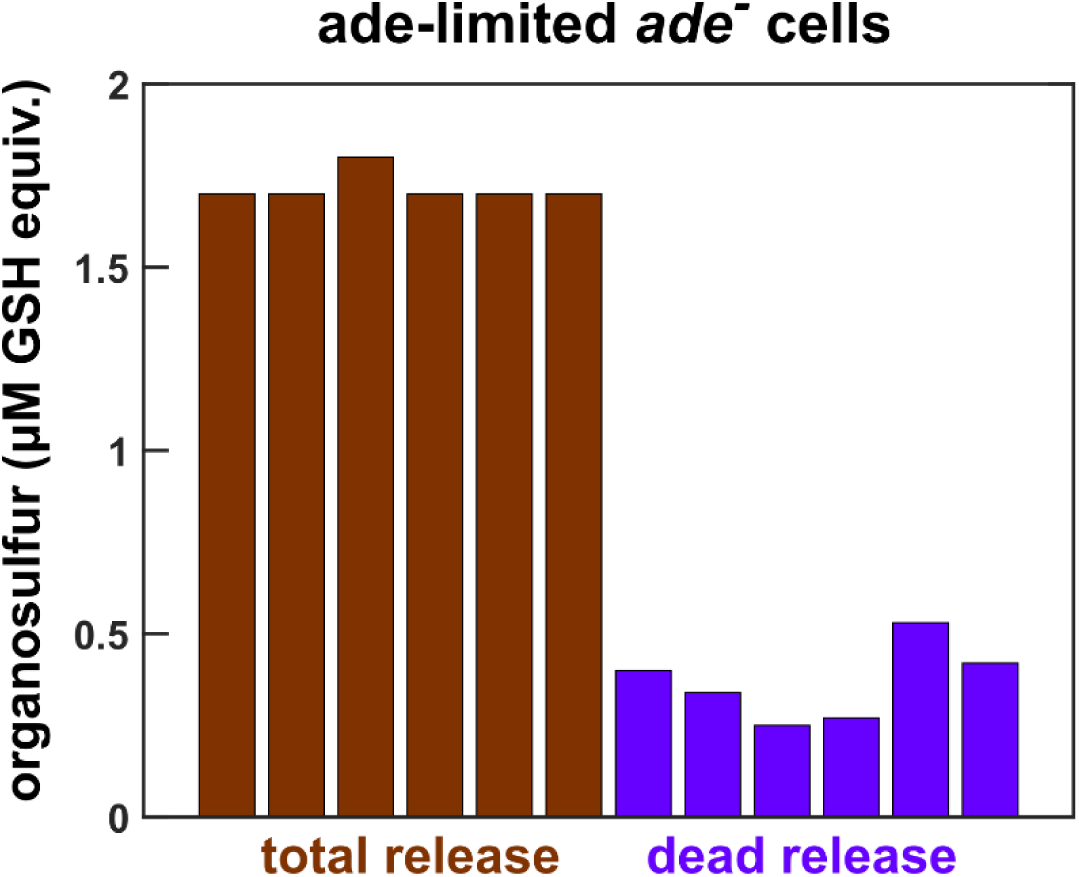
Live adenine-limited *ade*^*-*^ cells release organosulfurs. *ade*^-^cells (WY1340; WY1598) were grown in adenine-limited chemostats at 8-hr doubling time. After cultures had reached steady-state cell density (71∼73 hrs), a sample was taken to assay for total cell density, live cell density, and dead cell density using flow cytometry. Simultaneously, another sample was filtered to harvest supernatant, and a third sample was taken to harvest cells to make extracts. Total organosulfurs in supernatants (brown) and cell extracts were measured via the *met10*-based bioassay. From organosulfur concentrations in extracts and the total number of cells harvested, we calculated fmole organosulfurs per cell. We inferred the contribution to total release by dead cells by multiplying dead cell density with fmole organosulfur/cell.

**Fig S9.**
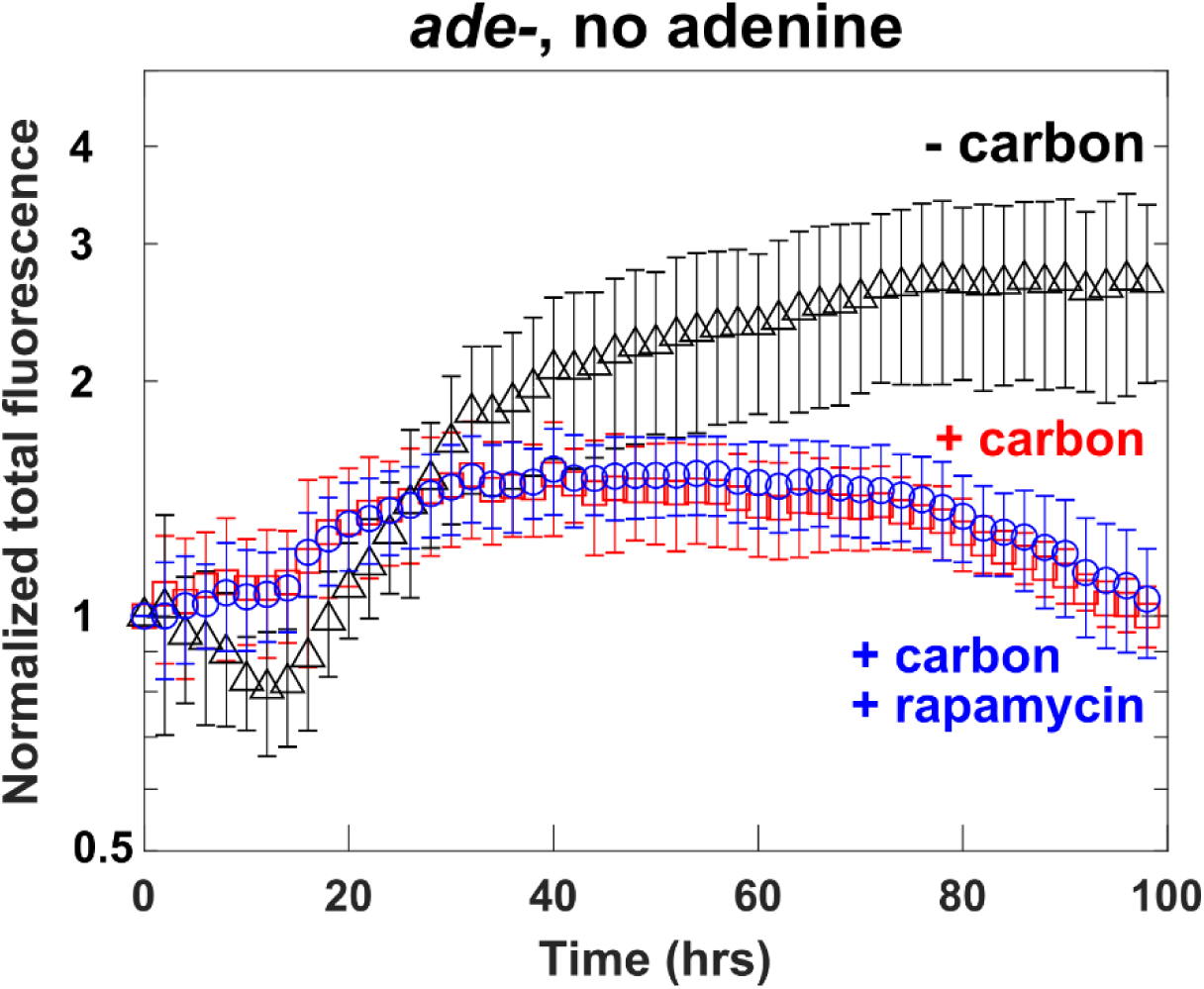
Adenine-limited *ade*^*-*^ cells display aspects of nutrient-growth dysregulation. Multiple lines of evidence suggest that adenine-limited *ade*^-^cells suffer nutrient-growth dysregulation. In our previous work, we found that in very low concentrations (0.1∼0.2 µM) of hypoxanthine (which can be converted to adenine by cells), *ade*^-^cells initially divided but then died at a faster rate than during adenine starvation, consistent with nutrient-growth dysregulation during adenine limitation (Fig 3C in (Hart et al., 2019a)). Here, exponentially growing *ade*^-^cells (WY1340) were washed in sterile water, and cultured in minimal medium lacking adenine, either with glucose (“+ carbon”, red) or without glucose (“-carbon”, black) for 24 hours before imaging in the same medium. For the rapamycin sample (“+carbon+rapamycin”, blue), rapamycin was added at the beginning of imaging. After an initial long lag, *ade*^-^cells died (red). Similar to *lys*^-^cells, cell death during adenine starvation was mitigated by removing glucose (compare red with black). Unlike *lys*^-^cells, cell death was not inhibited by rapamycin (compare red with blue). This suggests that during purine starvation, although TORC1 is already inactivated, other pathways (such as Ras/PKA) are still active, contributing to nutrient-growth dysregulation. Error bars correspond to two standard deviations for 6 replicate wells.

**Fig S10.**
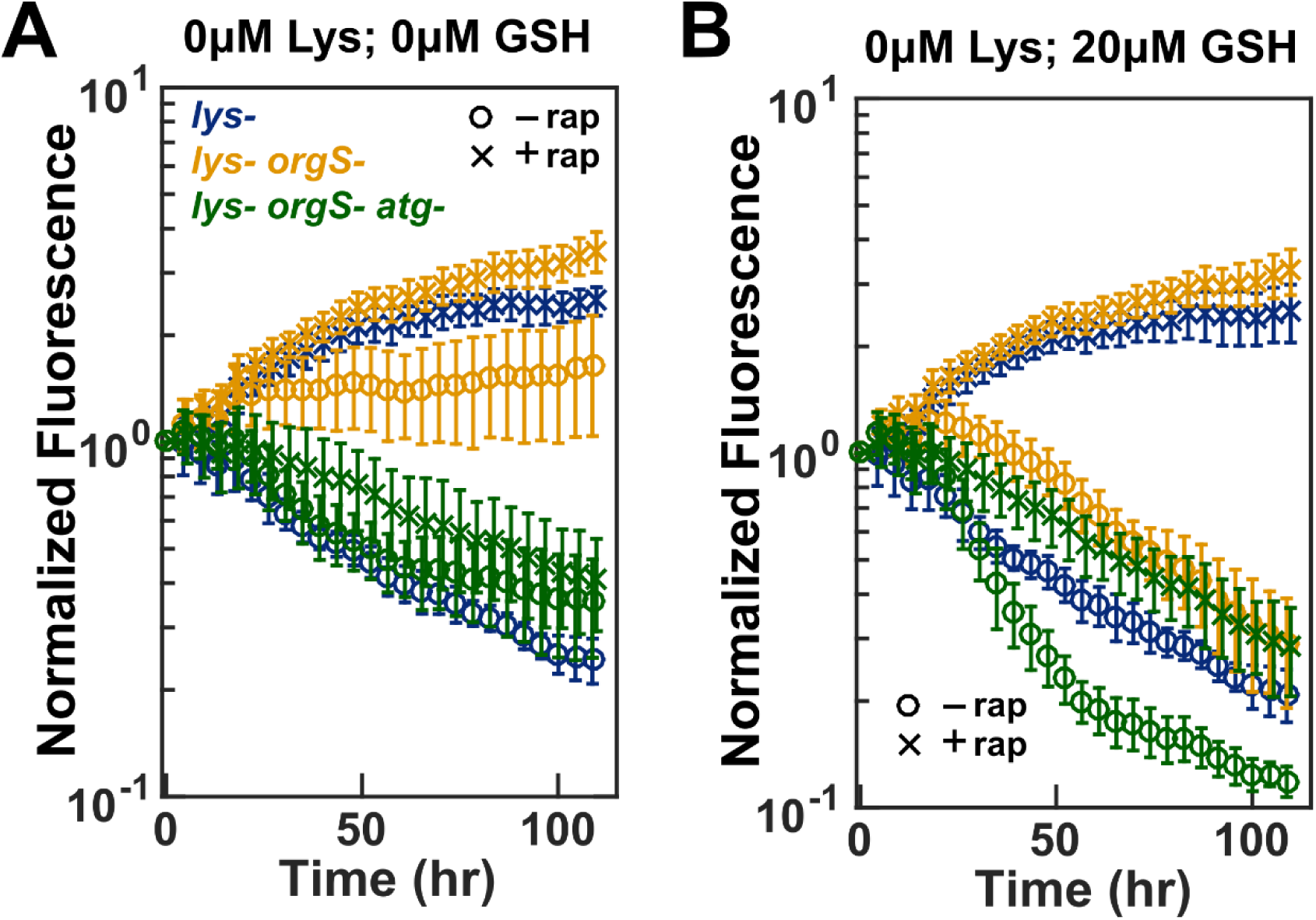
*lys*^-^*orgS*^-^survives lysine limitation better than *lys*^-^if organosulfur is also limited. *lys*^-^(WY2429, blue), *lys*^-^*orgS*^-^(WY1604, orange), and *lys*^-^*orgS*^-^*atg5*^-^(WY2370, green) were grown to exponential phase in SD supplemented with excess lysine (164 µM) and excess GSH (134 µM). These cells were washed and starved in SD without lysine or GSH for 5 hours prior to imaging in indicated conditions with 1 µM rapamycin (crosses) or without rapamycin (circles). Total fluorescence normalized against time zero are plotted. (**A**) When GSH was limited, *lys*^-^*orgS*^-^survived better than *lys*^-^(orange circles above blue circles). TORC1 inhibition by rapamycin improved the survival of both *lys*^-^*orgS*^-^and *lys*^-^to comparable levels (orange and blue crosses). *lys*^-^*orgS*^-^*atg5*^-^survived poorly even when TORC1 was shut down (green crosses). (**B**) At high GSH, both *lys*^-^*orgS*^-^and *lys*^-^survived poorly (orange circles trending down at a similar slope as blue circles), and this poor viability was rescued by rapamycin (blue and orange crosses). *lys*^-^*orgS*^-^*atg5*^-^survived poorly in the presence or absence of rapamycin (green). The ∼2-fold initial increase in the top curves was due to cell swelling (Hart et al., 2019b). Error bars represent two SEM (standard error of mean) from six wells.

**Fig S11.**
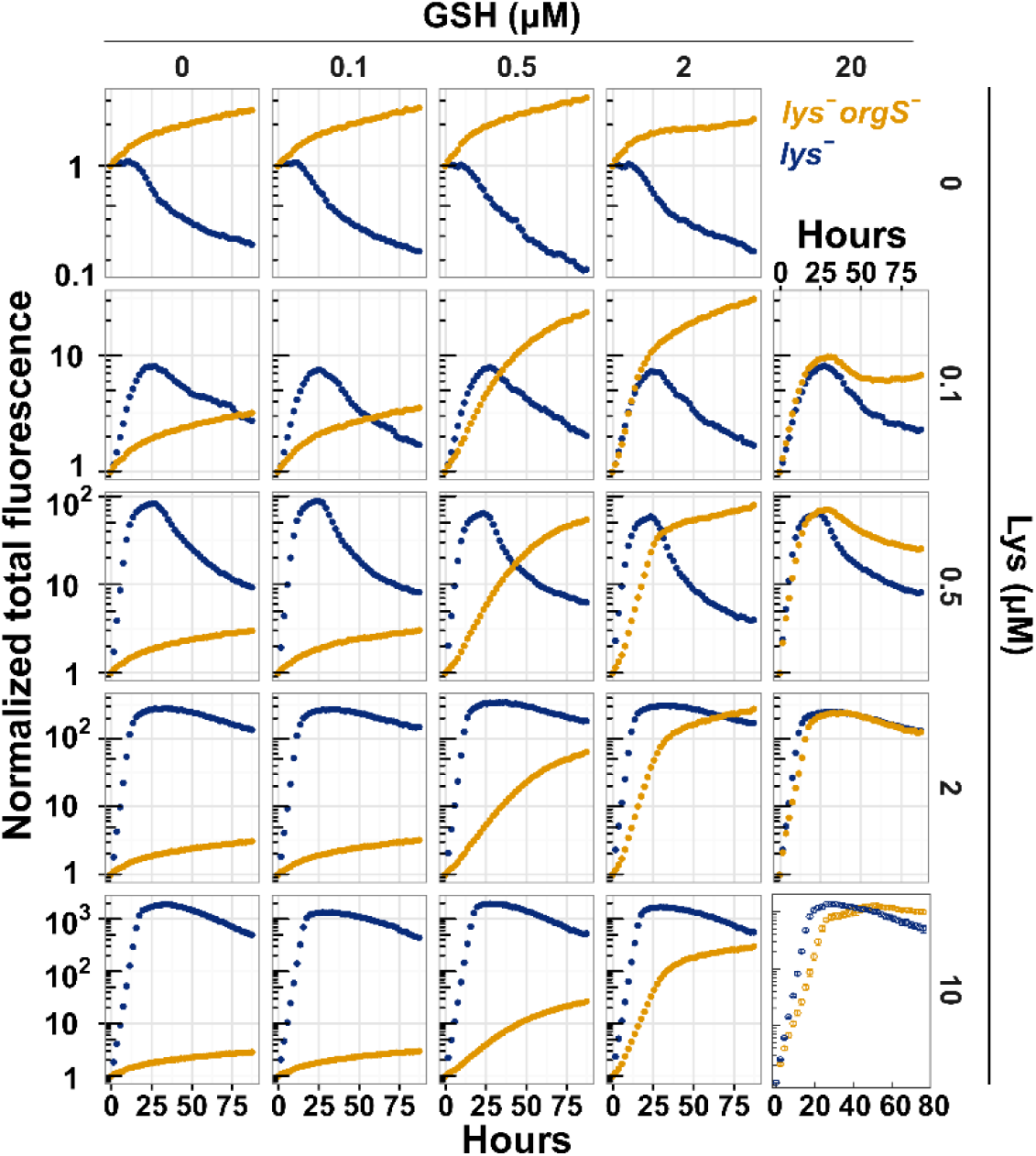
Comparing the growth and death profiles of *lys*^-^*orgS*^-^and *lys*^-^cells. *lys*^-^(WY2429, blue) and *lys*^-^*orgS*^-^(WY1604, orange) cells were grown in SD+ excess lysine (164 µM) and excess GSH (134 µM) to exponential phase. These cells were washed and starved in SD for 24 hours and imaged in various concentrations of GSH and lysine. During the growth phase, *lys*^-^grew faster than *lys*^-^*orgS*^-^. After lysine was exhausted, *lys*^-^*orgS*^-^survived better than *lys*^-^. Note that in the microscopy assay, the minimal medium did not contain GSX or other excreted compounds found in culture supernatants, and thus results are not directly comparable to competition experiments. Regardless, and consistent with the competition experiment, *lys*^-^*orgS*^-^cells grew faster than *lys*^-^cells under certain conditions. In this experiment, at 2 µM GSH and 0.1 µM lysine, the maximal growth rate achieved by *lys*^-^*orgS*^-^was 0.153 ± 0.009/hr (mean ± 2 standard error of mean), greater than the 0.133 ± 0.004/hr achieved by *lys*^-^. Fluorescence intensities of various time points are normalized against that at time zero.

**Fig S12.**
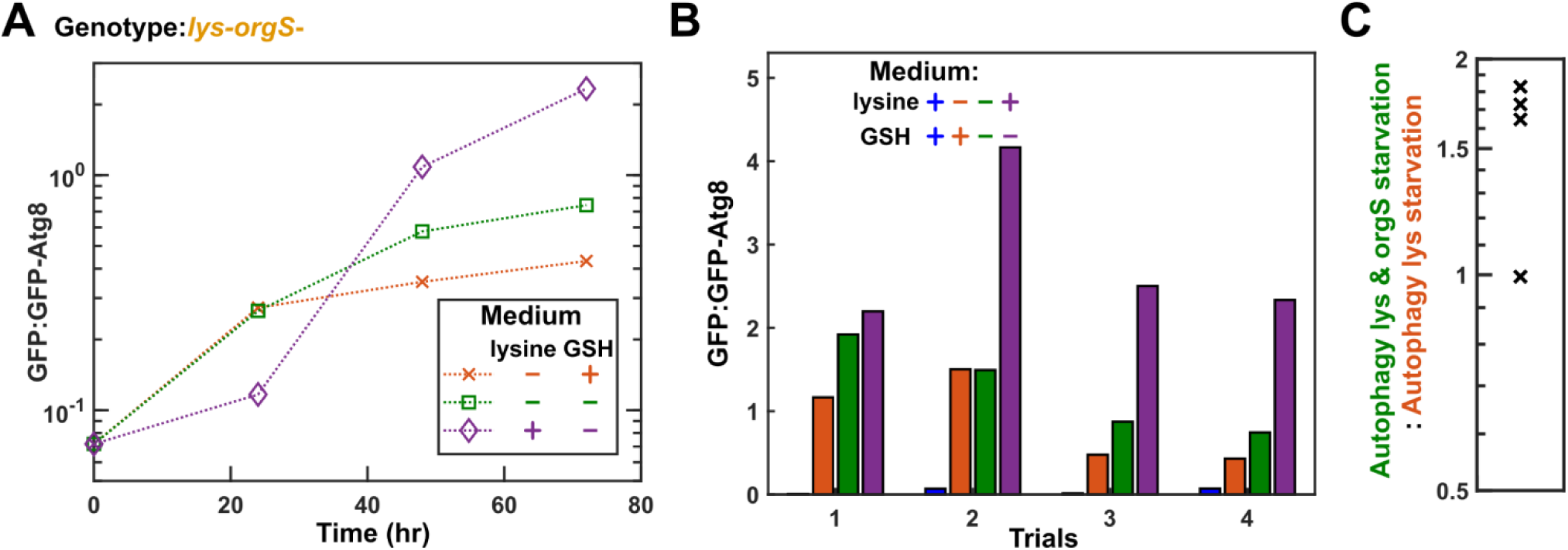
Autophagy activity is higher when *lys*^-^*orgS*^-^cells are starved of both organosulfurs and lysine compared to starved of lysine alone. The ideal comparison of autophagy between *lys*^*-*^ and *lys*^*-*^*orgS*^*-*^ cells has multiple technical difficulties. First, cells should ideally be cultured in an environment that mimics the original evolutionary environment (i.e. in low concentrations of lysine and organosulfurs), and then the steady-state autophagy activities can be measured. This means that *lys*^*-*^*orgS*^*-*^ cells would need to be cultured in a chemostat dual-limited for sulfur and lysine. However, the theory of chemostat is based on single nutrient limitation (Novick and Szilard, 1951), and ensuring dual limitation is nontrivial. Thus, we assayed autophagy in batch starvation cultures. Second, it can be difficult to compare autophagy activities between different genotypes. For example, the kinetics of autophagy and death differed drastically between *lys*^*-*^ and *lys*^*-*^*orgS*^*-*^. When supplied with excess lysine and no organosulfurs, *lys*^*-*^*orgS*^*-*^ cells continued to divide for multiple rounds using internal organosulfur storage, and autophagy induction was very slow. In contrast, lysine-starved *lys*^*-*^ cells died quickly. Thus, a comparison between the two genotypes is difficult. For these reasons, we compared autophagy activities in batch cultures of a single genotype (*lys*^*-*^*orgS*^*-*^; WY2520) under different starvation conditions. Using the GFP-Atg8 cleavage assay, we observed a moderate but significant increase in autophagy when *lys*^*-*^*orgS*^*-*^ cells were starved for both lysine and organosulfurs as opposed to only lysine starvation. (**A**) Time course of autophagy induction during lysine starvation (orange), organosulfur starvation (purple), and dual starvation (green). (**B**) Results from four trials (the fourth trial is identical to A). Autophagy of exponential cultures (first bar in each set) and of singly or doubly starved cultures at 72 hrs (second to fourth bars in each set) are plotted. (**C**) Autophagy activity was higher in cells starved for both lysine and organosulfurs than in cells starved for lysine only (P=0.03, one-tailed one-sample t test).

**Fig S13.**
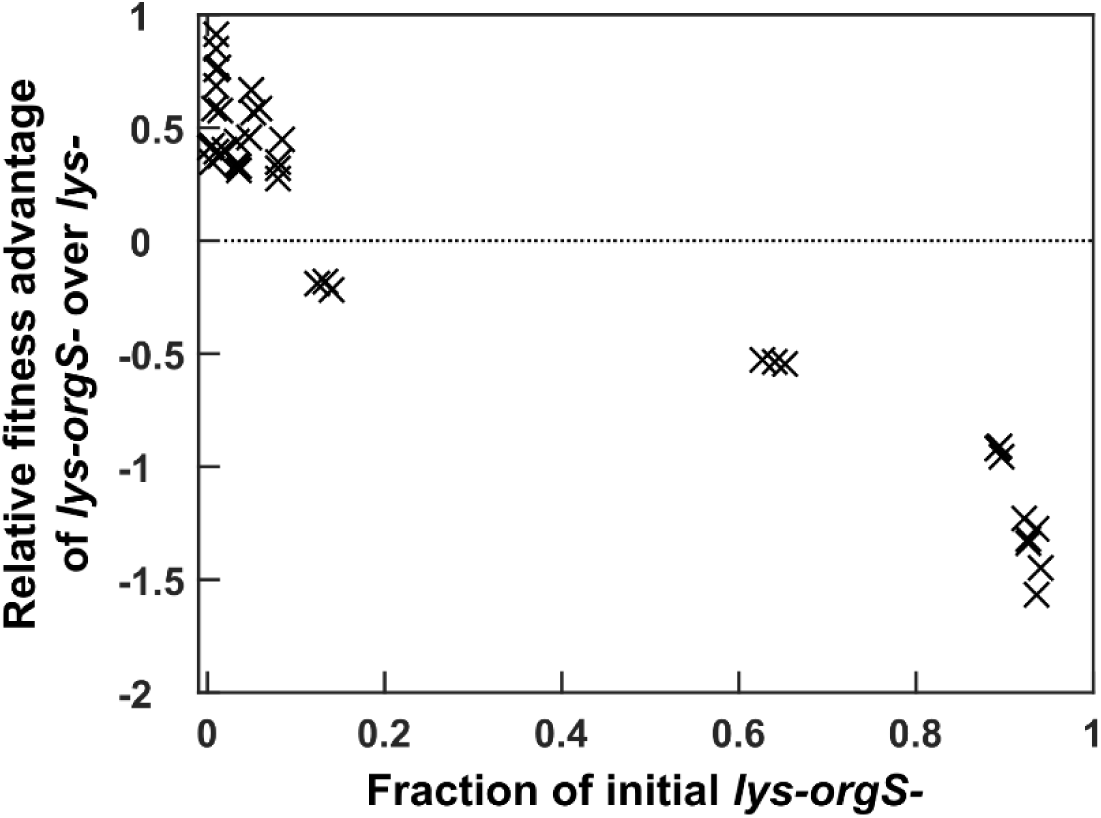
Negative frequency-dependent selection for *lys*^-^*orgS*^-^. BFP-tagged *lys*^-^*orgS*^-^(WY2072 or WY2073) and mCherry-tagged *lys*^-^(WY2039 or WY2045) were competed in a lysine-limited environment by coculturing with a lysine-releasing strain (WY1340). Strain ratios over time were measured by flow cytometry (Fig 6C). For each trajectory, we computed the slope of ln(*lys*^-^*orgS*^-^: *lys*^-^) over three consecutive time points, and chose the steepest slope. Since our time unit was generation, we divided this slope (/generation) by ln2/generation and obtained a dimensionless number representing the relative fitness difference between *lys*^-^*orgS*^-^and *lys*^-^. We then plotted the relative fitness difference against the fraction of *lys*^-^*orgS*^-^at the beginning of the time window used to calculate the steepest slope. Dotted line marks equal fitness between the two strains. The fitness advantage of *lys*^-^*orgS*^-^over *lys*^-^decreases as the fraction of *lys*^-^*orgS*^-^increases (i.e. negative frequency-dependent).

**Fig S14.**
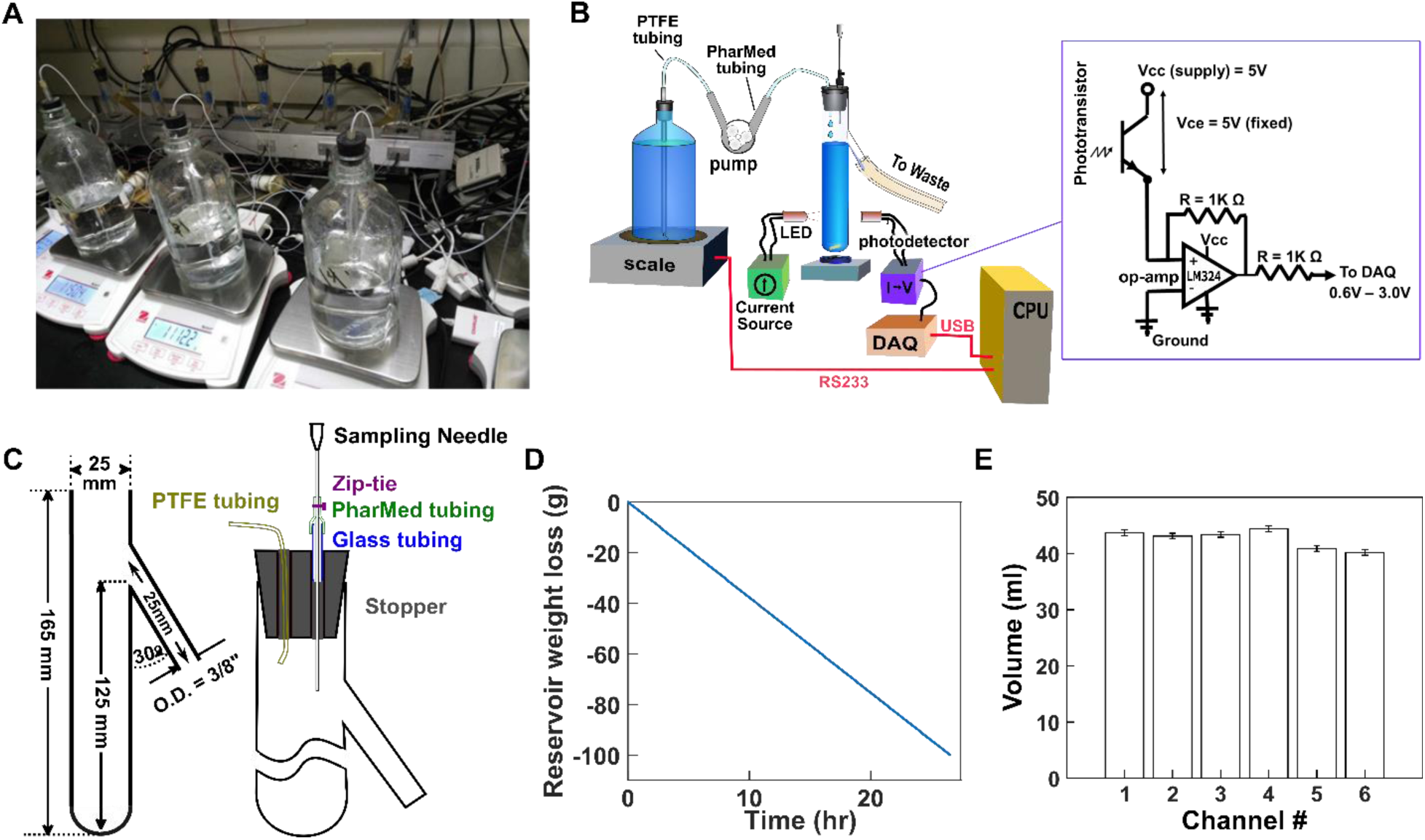
Continuous culturing device. The continuous culturing device (**A**) consists of six channels which can independently operate as a chemostat or a turbidostat. Each channel (**B**) consists of a culturing vessel (center), a magnetic stirrer, an LED-phototransistor optical detector for OD measurement, a computer activated pump, a media reservoir (left), and a scale for measuring media reservoir (and flow rate). A LabView program running on the CPU uses data from the scale or the optical detector to control the pump, which maintains a constant average OD in the turbidostat mode, or a constant average flow rate in the chemostat mode. Each vessel (**C**) consists of a Pyrex test tubes modified by adding a waste outlet of adequate diameter and slope to ensure a reliable flow of waste driven by gravity. The vessel’s rubber stopper had a sampling port consisting of a needle that can be raised and lowered through a segment of PharMed tubing which was held in place by glass tubing inserted into the stopper. The tightness of the seal between the sampling needle and PharMed tubing can be adjusted using a Zip-tie, allowing easy motion of the needle, while maintaining its position when stationary. The six vessels, stirrers, and photodetectors are held in position in a frame cut from an aluminum bar. The signal from each phototransistor was converted to a voltage using an op-amp current to voltage converter (box in **B**). (**D**) Constant flow rates in chemostats. An example is shown. (**E**) Culturing vessel volume averages 43 ml. Individual culturing vessel volume was used for converting doubling time to flow rate, and was measured to ∼0.5 ml resolution (limited by minimum outflow drop size; error bar).

**Supplementary Text 1**

Although some biochemistry results seemed to conflict with the nutrient-growth regulation model (Fig 1), we can offer alternative explanations for these results. For example, during unnatural limitation of leucine, histidine, or lysine, TORC1 seemed to be inactivated toward the one or two tested substrates (Binda et al., 2009; Kamada, 2017). We argue that TORC1 could be active against other substrates. Consistent with this notion, inactivating TORC1 via rapamycin rescued poor viability of *lys*^*-*^ cells during lysine starvation (Fig 2A). In addition, the Ras/PKA pathway could also be active, contributing to nutrient-growth dysregulation. As another example, leucine-sensing mechanisms have been found in yeast (González and Hall, 2017; Zhang et al., 2018), suggesting that leucine limitation may be interpreted as a natural limitation. Regardless of the leucine-sensing mechanism, *leu*^*-*^ cells still suffered poor viability during leucine starvation compared to during natural starvation (Boer et al., 2008; Gresham et al., 2011; Petti et al., 2011). One possibility is that growth inhibition by leucine starvation is overwhelmed by growth promotion by natural nutrients.

We argue that when different assays generate conflicting results, fitness phenotypes (e.g. cell growth or death rate) should always be given the highest weight. This is because 1) fitness is the final read-out of all the known and the unknown biochemistry inside and outside of a cell, and 2) natural selection acts on fitness.

**Table S1. Strains used in this study**.

**Table S2. Mutations in evolved clones**

Table S2: All clones with the alias “ACI” are from chemostat evolution experiment. All other clones are from coculture evolution experiments, a fraction of which (i.e. “CT” strains) are from an earlier study (Waite and Shou, 2012). Grey shading: auxotrophic strain. For WY2467, we identified it as *orgS-* from growth patterns. However, its genetic basis is unclear since among identified mutations, none is known to affect organosulfur biosynthesis. Matched color shading: mutation adaptive to lysine limitation shared between a *lys*^*-*^ clone and a *lys*^*-*^*orgS*^*-*^ clone from the same culture.

## References

Bachhawat AK, Thakur A, Kaur J, Zulkifli M. 2013. Glutathione transporters. Biochim Biophys Acta BBA - Gen Subj, Cellular functions of glutathione 1830:3154–3164. doi: 10.1016/j.bbagen.2012.11.018

Basan M, Hui S, Okano H, Zhang Z, Shen Y, Williamson JR, Hwa T. 2015. Overflow metabolism in Escherichia coli results from efficient proteome allocation. Nature 528:99–104. doi: 10.1038/nature15765

Beliaev AS, Romine MF, Serres M, Bernstein HC, Linggi BE, Markillie LM, Isern NG, Chrisler WB, Kucek LA, Hill EA, Pinchuk GE, Bryant DA, Steven Wiley H, Fredrickson JK, Konopka A. 2014. Inference of interactions in cyanobacterial–heterotrophic co-cultures via transcriptome sequencing. ISME J 8:2243–2255. doi: 10.1038/ismej.2014.69

Binda M, Péli-Gulli M-P, Bonfils G, Panchaud N, Urban J, Sturgill TW, Loewith R, De Virgilio C. 2009. The Vam6 GEF controls TORC1 by activating the EGO complex. Mol Cell 35:563–573. doi: 10.1016/j.molcel.2009.06.033

Boer VM, Amini S, Botstein D. 2008. Influence of genotype and nutrition on survival and metabolism of starving yeast. Proc Natl Acad Sci 105:6930–6935. doi: 10.1073/pnas.0802601105

Boer VM, Crutchfield CA, Bradley PH, Botstein D, Rabinowitz JD. 2010. Growth-limiting intracellular metabolites in yeast growing under diverse nutrient limitations. Mol Biol Cell 21:198–211. doi: 10.1091/mbc.E09-07-0597

Brauer MJ, Huttenhower C, Airoldi EM, Rosenstein R, Matese JC, Gresham D, Boer VM, Troyanskaya OG, Botstein D. 2008. Coordination of Growth Rate, Cell Cycle, Stress Response, and Metabolic Activity in Yeast. Mol Biol Cell 19:352–367. doi: 10.1091/mbc.E07-08-0779

Campbell K, Vowinckel J, Muelleder M, Malmsheimer S, Lawrence N, Calvani E, Miller-Fleming L, Alam MT, Christen S, Keller MA, Ralser M. 2015. Self-establishing communities enable cooperative metabolite exchange in a eukaryote. eLife e09943. doi: 10.7554/eLife.09943

Carini P, Campbell EO, Morré J, Sañudo-Wilhelmy SA, Cameron Thrash J, Bennett SE, Temperton B, Begley T, Giovannoni SJ. 2014. Discovery of a SAR11 growth requirement for thiamin’s pyrimidine precursor and its distribution in the Sargasso Sea. ISME J 8:1727–1738. doi: 10.1038/ismej.2014.61

Castrillo JI, Zeef LA, Hoyle DC, Zhang N, Hayes A, Gardner DC, Cornell MJ, Petty J, Hakes L, Wardleworth L. 2007. Growth control of the eukaryote cell: a systems biology study in yeast. J Biol 6:4.

Conrad M, Schothorst J, Kankipati HN, Zeebroeck GV, Rubio-Texeira M, Thevelein JM. 2014. Nutrient sensing and signaling in the yeast Saccharomyces cerevisiae. FEMS Microbiol Rev 38:254–299. doi: 10.1111/1574-6976.12065

Dhaoui M, Auchère F, Blaiseau P-L, Lesuisse E, Landoulsi A, Camadro J-M, Haguenauer-Tsapis R, Belgareh-Touzé N. 2011. Gex1 is a yeast glutathione exchanger that interferes with pH and redox homeostasis. Mol Biol Cell 22:2054–2067. doi: 10.1091/mbc.E10-11-0906

D’Souza G, Kost C. 2016. Experimental Evolution of Metabolic Dependency in Bacteria. PLOS Genet 12:e1006364. doi: 10.1371/journal.pgen.1006364

D’Souza G, Waschina S, Pande S, Bohl K, Kaleta C, Kost C. 2014. Less Is More: Selective Advantages Can Explain the Prevalent Loss of Biosynthetic Genes in Bacteria. Evolution 68:2559–2570. doi: 10.1111/evo.12468

Dykhuizen D. 1978. Selection for Tryptophan Auxotrophs of Escherichia coli in Glucose-Limited Chemostats as a Test of the Energy Conservation Hypothesis of Evolution. Evolution 32:125–150. doi: 10.2307/2407415

Gasch AP, Spellman PT, Kao CM, Carmel-Harel O, Eisen MB, Storz G, Botstein D, Brown PO. 2000. Genomic Expression Programs in the Response of Yeast Cells to Environmental Changes. Mol Biol Cell 11:4241–4257.

González A, Hall MN. 2017. Nutrient sensing and TOR signaling in yeast and mammals. EMBO J 36:397–408. doi: 10.15252/embj.201696010

Grant CM, MacIver FH, Dawes IW. 1996. Glutathione is an essential metabolite required for resistance to oxidative stress in the yeastSaccharomyces cerevisiae. Curr Genet 29:511–515. doi: 10.1007/BF02426954

Gresham D, Boer VM, Caudy A, Ziv N, Brandt NJ, Storey JD, Botstein D. 2011. System-Level Analysis of Genes and Functions Affecting Survival During Nutrient Starvation in Saccharomyces cerevisiae. Genetics 187:299–317. doi: 10.1534/genetics.110.120766

Gresham D, Dunham MJ. 2014. The enduring utility of continuous culturing in experimental evolution. Genomics, Experimental evolution and the use of genomics 104:399–405. doi: 10.1016/j.ygeno.2014.09.015

Guthrie C, Fink GR. 1991. Guide to yeast genetics and molecular biology. Academic Press.

Harcombe W. 2010. Novel cooperation experimentally evolved between species. Evolution 64:2166–2172. doi: 10.1111/j.1558-5646.2010.00959.x

Harcombe WR, Chacón JM, Adamowicz EM, Chubiz LM, Marx CJ. 2018. Evolution of bidirectional costly mutualism from byproduct consumption. Proc Natl Acad Sci 115:12000–12004.

Hart & Pineda, Chen, Chichun, Green, Robin, Shou W, Shou W. 2019. Disentangling strictly self-serving mutations from win-win mutations in a mutualistic microbial community. ELife Accept.

Hart SFM, Mi H, Green R, Xie L, Pineda JMB, Momeni B, Shou W. 2019a. Uncovering and resolving challenges of quantitative modeling in a simplified community of interacting cells. PLOS Biol 17:e3000135. doi: 10.1371/journal.pbio.3000135

Hart SFM, Skelding D, Waite AJ, Burton JC, Shou W. 2019b. High-throughput quantification of microbial birth and death dynamics using fluorescence microscopy. Quant Biol. doi: 10.1007/s40484-018-0160-7

Helliwell KE, Wheeler GL, Leptos KC, Goldstein RE, Smith AG. 2011. Insights into the Evolution of Vitamin B12 Auxotrophy from Sequenced Algal Genomes. Mol Biol Evol 28:2921–2933. doi: 10.1093/molbev/msr124

Hess DC, Lu W, Rabinowitz JD, Botstein D. 2006. Ammonium Toxicity and Potassium Limitation in Yeast. PLoS Biol 4:e351. doi: 10.1371/journal.pbio.0040351

Hillesland Kristina Linnea. 2017. Evolution on the bright side of life: microorganisms and the evolution of mutualism. Ann N Y Acad Sci 0. doi: 10.1111/nyas.13515

Hu J, Fan L, Chen Q, Dong Y. 2017. RNA-Seq-based transcriptomic and metabolomic analysis reveal stress responses and programmed cell death induced by acetic acid in *Saccharomyces cerevisiae*. Sci Rep 7:42659. doi: 10.1038/srep42659

Huang J, Klionsky DJ. 2007. Autophagy and Human Disease. Cell Cycle 6:1837–1849. doi: 10.4161/cc.6.15.4511

Huang W-P, Shintani T, Xie Z. 2014. Assays for Autophagy I: The Cvt Pathway and Nonselective Autophagy In: Xiao W, editor. Yeast Protocols, Methods in Molecular Biology. New York, NY: Springer. pp. 153–164. doi: 10.1007/978-1-4939-0799-1_10

Ishikawa T. 1992. The ATP-dependent glutathione S-conjugate export pump. Trends Biochem Sci 17:463–468.

Jewell JL, Guan K-L. 2013. Nutrient signaling to mTOR and cell growth. Trends Biochem Sci 38:233–242. doi: 10.1016/j.tibs.2013.01.004

Jiang X, Zerfaß C, Feng S, Eichmann R, Asally M, Schäfer P, Soyer OS. 2018. Impact of spatial organization on a novel auxotrophic interaction among soil microbes. ISME J 1. doi: 10.1038/s41396-018-0095-z

Kamada Y. 2017. Novel tRNA function in amino acid sensing of yeast Tor complex1. Genes Cells Devoted Mol Cell Mech 22:135–147. doi: 10.1111/gtc.12462

Kingsbury JM, Sen ND, Cardenas ME. 2015. Branched-Chain Aminotransferases Control TORC1 Signaling in Saccharomyces cerevisiae. PLoS Genet 11. doi: 10.1371/journal.pgen.1005714

Kinnersley MA, Holben WE, Rosenzweig F. 2009. E Unibus Plurum: genomic analysis of an experimentally evolved polymorphism in Escherichia coli. PLoS Genet 5:e1000713. doi: 10.1371/journal.pgen.1000713

Klosinska MM, Crutchfield CA, Bradley PH, Rabinowitz JD, Broach JR. 2011. Yeast cells can access distinct quiescent states. Genes Dev 25:336–349. doi: 10.1101/gad.2011311

Laland KN, Odling-Smee FJ, Feldman MW. 1999. Evolutionary consequences of niche construction and their implications for ecology. Proc Natl Acad Sci 96:10242–10247. doi: 10.1073/pnas.96.18.10242

Laxman S, Sutter BM, Tu BP. 2014. Methionine is a signal of amino acid sufficiency that inhibits autophagy through the methylation of PP2A. Autophagy 10:386–387. doi: 10.4161/auto.27485

Lin CH, MacGurn JA, Chu T, Stefan CJ, Emr SD. 2008. Arrestin-Related Ubiquitin-Ligase Adaptors Regulate Endocytosis and Protein Turnover at the Cell Surface. Cell 135:714–725. doi: 10.1016/j.cell.2008.09.025

Masselot M, Robichon-Szulmajster H. 1975. Methionine biosynthesis in Saccharomyces cerevisiae: I. Genetical analysis of auxotrophic mutants. MGG Mol Gen Genet 139. doi: 10.1007/BF00264692

Matsumoto K, Uno I, Ishikawa T. 1983. Control of cell division in Saccharomyces cerevisiae mutants defective in adenylate cyclase and cAMP-dependent protein kinase. Exp Cell Res 146:151–161. doi: 10.1016/0014-4827(83)90333-6

Mee MT, Collins JJ, Church GM, Wang HH. 2014. Syntrophic exchange in synthetic microbial communities. Proc Natl Acad Sci 201405641. doi: 10.1073/pnas.1405641111

Melamud E, Vastag L, Rabinowitz JD. 2010. Metabolomic Analysis and Visualization Engine for LC-MS Data. Anal Chem 82:9818–9826. doi: 10.1021/ac1021166

Momeni B, Waite AJ, Shou W. 2013. Spatial self-organization favors heterotypic cooperation over cheating. eLife 2:e00960. doi: 10.7554/eLife.00960

Morris JJ, Lenski RE, Zinser ER. 2012. The Black Queen Hypothesis: evolution of dependencies through adaptive gene loss. mBio 3. doi: 10.1128/mBio.00036-12

Novick A, Szilard L. 1951. Experiments on spontaneous and chemically induced mutations of bacteria growing in the Chemostat. Cold Spring Harb Symp Quant Biol 16:337–43.

Paczia N, Nilgen A, Lehmann T, Gätgens J, Wiechert W, Noack S. 2012. Extensive exometabolome analysis reveals extended overflow metabolism in various microorganisms. Microb Cell Factories 11:122. doi: 10.1186/1475-2859-11-122

Pande S, Kaftan F, Lang S, Svatoš A, Germerodt S, Kost C. 2016. Privatization of cooperative benefits stabilizes mutualistic cross-feeding interactions in spatially structured environments. ISME J 10:1413–1423. doi: 10.1038/ismej.2015.212

Petti, Crutchfield CA, Rabinowitz JD. 2011. Survival of starving yeast is correlated with oxidative stress response and nonrespiratory mitochondrial function. Proc Natl Acad Sci 108:E1089–E1098.

Ponomarova O, Gabrielli N, Sévin DC, Mülleder M, Zirngibl K, Bulyha K, Andrejev S, Kafkia E, Typas A, Sauer U. 2017. Yeast Creates a Niche for Symbiotic Lactic Acid Bacteria through Nitrogen Overflow. Cell Syst 5:345–357.

Pópulo H, Lopes JM, Soares P. 2012. The mTOR Signalling Pathway in Human Cancer. Int J Mol Sci 13:1886–1918. doi: 10.3390/ijms13021886

Rebbeor JF, Connolly GC, Dumont ME, Ballatori N. 1998. ATP-dependent transport of reduced glutathione in yeast secretory vesicles. Biochem J 334 (Pt 3):723–729.

Regenberg B, Grotkjær T, Winther O, Fausbøll A, Åkesson M, Bro C, Hansen LK, Brunak S, Nielsen J. 2006. Growth-rate regulated genes have profound impact on interpretation of transcriptome profiling in Saccharomyces cerevisiae. Genome Biol 7:R107. doi: 10.1186/gb-2006-7-11-r107

Rodionova IA, Li X, Plymale AE, Motamedchaboki K, Konopka AE, Romine MF, Fredrickson JK, Osterman AL, Rodionov DA. 2015. Genomic distribution of B-vitamin auxotrophy and uptake transporters in environmental bacteria from the C hloroflexi phylum. Environ Microbiol Rep 7:204–210.

Rosebrock AP, Caudy AA. 2017. Metabolite Extraction from Saccharomyces cerevisiae for Liquid Chromatography–Mass Spectrometry. Cold Spring Harb Protoc 2017:pdb.prot089086. doi: 10.1101/pdb.prot089086

Saldanha AJ, Brauer MJ, Botstein D. 2004. Nutritional Homeostasis in Batch and Steady-State Culture of Yeast. Mol Biol Cell 15:4089–4104. doi: 10.1091/mbc.E04-04-0306

Shou W, Ram S, Vilar JM. 2007. Synthetic cooperation in engineered yeast populations. Proc Natl Acad Sci USA 104:1877–1882. doi: 10.1073/pnas.0610575104

Skelding D, Hart SF, Vidyasagar T, Pozhitkov AE, Shou W. 2018. Developing a low-cost milliliter-scale chemostat array for precise control of cellular growth. Quant Biol 6:129–141.

Slavov N, Botstein D. 2013. Decoupling nutrient signaling from growth rate causes aerobic glycolysis and deregulation of cell size and gene expression. Mol Biol Cell 24:157–168. doi: 10.1091/mbc.E12-09-0670

Slavov N, Botstein D. 2011. Coupling among growth rate response, metabolic cycle, and cell division cycle in yeast. Mol Biol Cell 22:1997–2009. doi: 10.1091/mbc.E11-02-0132

Stams AJM, Bok FAM de, Plugge CM, Eekert MHA van, Dolfing J, Schraa G. 2006. Exocellular electron transfer in anaerobic microbial communities. Environ Microbiol 8:371–382. doi: 10.1111/j.1462-2920.2006.00989.x

Sutter BM, Wu X, Laxman S, Tu BP. 2013. Methionine Inhibits Autophagy and Promotes Growth by Inducing the SAM-Responsive Methylation of PP2A. Cell 154:403–415. doi: 10.1016/j.cell.2013.06.041

Szenk M, Dill KA, de Graff AMR. 2017. Why Do Fast-Growing Bacteria Enter Overflow Metabolism? Testing the Membrane Real Estate Hypothesis. Cell Syst 5:95–104. doi: 10.1016/j.cels.2017.06.005

Takeshige K, Baba M, Tsuboi S, Noda T, Ohsumi Y. 1992. Autophagy in yeast demonstrated with proteinase-deficient mutants and conditions for its induction. J Cell Biol 119:301–311. doi: 10.1083/jcb.119.2.301

Thevelein JM, Cauwenberg L, Colombo S, De Winde JH, Donation M, Dumortier F, Kraakman L, Lemaire K, Ma P, Nauwelaers D, Rolland F, Teunissen A, Van Dijck P, Versele M, Wera S, Winderickx J. 2000. Nutrient-induced signal transduction through the protein kinase A pathway and its role in the control of metabolism, stress resistance, and growth in yeast. Enzyme Microb Technol 26:819–825. doi: 10.1016/S0141-0229(00)00177-0

Thomas D, Surdin-Kerjan Y. 1997. Metabolism of sulfur amino acids in Saccharomyces cerevisiae. Microbiol Mol Biol Rev 61:503–532.

Toda T, Cameron S, Sass P, Zoller M, Scott JD, McMullen B, Hurwitz M, Krebs EG, Wigler M. 1987. Cloning and characterization of BCY1, a locus encoding a regulatory subunit of the cyclic AMP-dependent protein kinase in Saccharomyces cerevisiae. Mol Cell Biol 7:1371–1377. doi: 10.1128/MCB.7.4.1371

Torggler R, Papinski D, Kraft C. 2017. Assays to Monitor Autophagy in Saccharomyces cerevisiae. Cells 6. doi: 10.3390/cells6030023

Unger MW, Hartwell LH. 1976. Control of cell division in Saccharomyces cerevisiae by methionyl-tRNA. Proc Natl Acad Sci U S A 73:1664–1668.

Wach A, Brachat A, Pöhlmann R, Philippsen P. 1994. New heterologous modules for classical or PCR-based gene disruptions in Saccharomyces cerevisiae. Yeast 10:1793–1808. doi: 10.1002/yea.320101310

Waite AJ, Shou W. 2014. Constructing synthetic microbial communities to explore the ecology and evolution of symbiosis. Methods Mol Biol Clifton NJ 1151:27–38. doi: 10.1007/978-1-4939-0554-6_2

Waite AJ, Shou W. 2012. Adaptation to a new environment allows cooperators to purge cheaters stochastically. Proc Natl Acad Sci 109:19079–19086. doi: 10.1073/pnas.1210190109

Xie Z, Nair U, Klionsky DJ. 2008. Atg8 controls phagophore expansion during autophagosome formation. Mol Biol Cell 19:3290–3298. doi: 10.1091/mbc.e07-12-1292

Yi L, Li H, Sun L, Liu L, Zhang C, Xi Z. 2009. A Highly Sensitive Fluorescence Probe for Fast Thiol-Quantification Assay of Glutathione Reductase. Angew Chem Int Ed 48:4034–4037. doi: 10.1002/anie.200805693

Zaman S, Lippman SI, Schneper L, Slonim N, Broach JR. 2009. Glucose regulates transcription in yeast through a network of signaling pathways. Mol Syst Biol 5:245. doi: 10.1038/msb.2009.2

Zaman S, Lippman SI, Zhao X, Broach JR. 2008. How Saccharomyces responds to nutrients. Annu Rev Genet 42:27–81.

Zamenhof S, Eichhorn HH. 1967. Study of microbial evolution through loss of biosynthetic functions: establishment of “defective” mutants. Nature 216:456–458.

Zengler K, Zaramela LS. 2018. The social network of microorganisms — how auxotrophies shape complex communities. Nat Rev Microbiol 1. doi: 10.1038/s41579-018-0004-5

Zhang J, Du G-C, Zhang Y, Liao X-Y, Wang M, Li Y, Chen J. 2010. Glutathione Protects Lactobacillus sanfranciscensis against Freeze-Thawing, Freeze-Drying, and Cold Treatment. Appl Environ Microbiol 76:2989–2996. doi: 10.1128/AEM.00026-09

Zhang W, Du G, Zhou J, Chen J. 2018. Regulation of Sensing, Transportation, and Catabolism of Nitrogen Sources in Saccharomyces cerevisiae. Microbiol Mol Biol Rev 82. doi: 10.1128/MMBR.00040-17

